# The evolution of socially modulated dispersal

**DOI:** 10.64898/2026.01.01.697281

**Authors:** Chon I Kam, Graeme D. Ruxton, Andy Gardner

## Abstract

Dispersal plays a key role in ecology and evolutionary biology, including as a mechanism that reduces local competition. However, dispersers often suffer a fitness disadvantage relative to nondispersers, which may lead individuals to prefer that others disperse rather than dispersing themselves. Hence, if individuals can force their neighbours to disperse, they may be favoured to do so rather than voluntarily dispersing for the good of others. Besides forcing dispersal, individuals might also improve their neighbours’ ability to disperse. Here, we investigate the evolutionary drivers and consequences of socially modulated dispersal. We find that: (1) there are three possible stable evolutionary outcomes, involving only voluntary dispersal, only socially modulated dispersal, or coexistence of the two; (2) although social modulation that does not improve dispersal success can only be favoured if it is less costly than dispersal itself, social modulation that facilitates dispersal can be favoured even if it is highly expensive for the facilitator; (3) facilitated dispersal may, counterintuitively, lead to a reduced overall level of dispersal; and (4) social modulation can be either welcome or unwelcome from the perspective of the disperser. These predictions emphasise that the evolution of dispersal depends on individuals’ abilities to both disperse and modulate.

## Introduction

Dispersal plays a key role in many areas of ecology and evolutionary biology (Clobert et al., 2001; Dieckmann et al., 1999). It shapes the genetics and demography of populations, and is itself subject to natural selection (Lowe & McPeek, 2014). Although it was once believed that dispersal would only be favoured by natural selection if changing location was directly beneficial to the disperser (Cohen, 1967; Gadgil, 1971), a landmark theoretical analysis by Hamilton & May (1977) demonstrated that dispersal can be favoured by kin selection even if dispersers suffer a net direct cost, on account of the reduced competition enjoyed by their genetic relatives whom they leave behind.

Accordingly, nondispersers often have a fitness advantage over dispersers (Bonte et al., 2012), which may lead to all individuals preferring that others disperse rather than dispersing themselves. While dispersal may be beneficial to the group and to the dispersing individuals in terms of their indirect fitness, those individuals that do not disperse and thus avoid the costs of dispersal may be more successful in terms of their direct fitness (cf. Motro, 1982a;1982b; 1983; Starrfelt & Kokko, 2010). Hence, if individuals could force the dispersal of their neighbours (i.e. patch mates)—for example, through eviction from the social group (Cant et al., 2001; Maag et al., 2022; Sarno et al., 2003; Thompson et al., 2017)—they may be favoured to do so rather than voluntarily dispersing for the good of others. In addition to forcing dispersal, individuals could also facilitate their neighbours’ dispersal, improving their likelihood of success in reaching their destination. Indeed, in some taxa individuals might not be able to disperse independently even if they would be favoured to do so, while in others independently dispersing individuals might only be able to travel a short distance owing to difficulties of limited motility—a situation that appears to have prompted the socially coordinated dispersal of microbes such as myxobacteria and cellular slime moulds (Bonner, 2015; Kaiser, 2004; Xue et al., 2011). However, the evolution of socially modulated dispersal remains underexplored.

Here, we investigate the evolutionary drivers and consequences of socially modulated dispersal. Aiming for a generic treatment that is not contingent on the biological details of a specific system, we develop a mathematical model of dispersal in a saturated, patch-structured population—based upon Hamilton & May’s (1977) classic model—to investigate how kin selection drives the co-evolutionary dynamics of voluntary versus socially modulated dispersal. We consider “forced dispersal”, in which individuals are made to disperse by others, and “facilitated dispersal”, in which individuals are both made to disperse and also enjoy a reduced cost of dispersal. This yields the conditions under which each form of dispersal may be evolutionarily maintained—and when both forms of dispersal can coexist—in terms of the costs experienced by the social modulator and the disperser as well as the genetic relatedness among social partners. We compare our mathematical results with those of classic analyses and translate them into qualitative and quantitative comparative predictions as to how different taxa are expected to exhibit different dispersal regimes.

## Results

### Forced dispersal

We assume an infinite, patch-structured population in which each patch contains a large number of individuals. Individuals choose whether to voluntarily disperse, but some individuals that do not disperse voluntarily may still be forced to disperse by their neighbours. Dispersal carries a risk of mortality, and each surviving disperser is independently assigned to a randomly chosen destination patch. Following dispersal, both original inhabitants and surviving dispersers compete for reproductive opportunities within their patch; those that are successful produce a large number of clonal offspring, and then all adults die. This returns the population to the beginning of the lifecycle. (See supplementary material for a sexual reproduction version of the model, which yields exactly the same results.)

Following these assumptions, we can write the relative fitness of a focal individual as

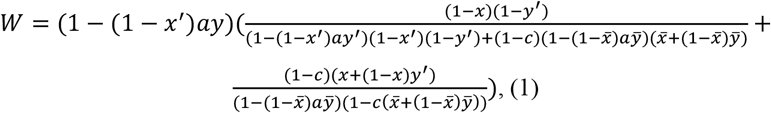

where *x* is the probability that she disperses voluntarily, *x’* is the average probability of voluntary dispersal among her neighbours, 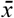 is the average probability of voluntary dispersal across the whole population, *y* is her proclivity for forcing neighbours to disperse, *y’* is the average proclivity for forcing among her neighbours, 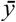 is the average proclivity for forcing across the population, *a* is the marginal cost—which we implement as reduced competitiveness for reproductive opportunities—of forcing a neighbour to disperse, and *c* is the probability that voluntary and forced dispersers die *en route* to their new patch (see supplementary material for detailed derivation, and Table 1 for a summary of model notation). We assume that an individual’s ability to force her neighbours to disperse is independent of whether she volunteers to disperse; relaxing this assumption leads to qualitatively similar results (see supplementary material for details). Fixing the levels of forced dispersal to zero (i.e. 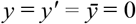) exactly recovers Hamilton & May’s (1977) model (see supplementary material for details).

**Table 1.** A summary of model notation.

| Symbol | Definition |
| --- | --- |
| $a$ | Cost of dispersal modulation |
| $c$ | Cost of dispersal |
| $f$ | Level of dispersal facilitation |
| $r$ | Relatedness between juvenile patch mates |
| $s$ | Degree of sibship among juvenile patch mates |
| $v$ | Relative reproductive value of an individual before dispersal |
| $W$ | Focal individual's relative fitness |
| $x$ | Focal individual's probability of voluntary dispersal |
| $x'$ | Average probability of voluntary dispersal in focal patch |
| $\bar{x}$ | Population average probability of voluntary dispersal |
| $x^*$ | Convergence stable probability of voluntary dispersal |
| $y$ | Focal individual's proclivity for modulating the dispersal of patch mates |
| $y'$ | Average proclivity for modulating dispersal in focal patch |
| $\bar{y}$ | Population average proclivity for modulating dispersal |
| $y^*$ | Convergence stable level of dispersal modulation |

Applying Taylor & Frank’s (1996) kin selection methodology, we find that natural selection favours an increase in voluntary dispersal if

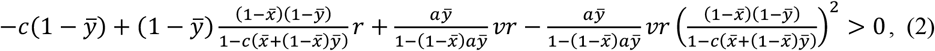

where 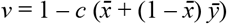 is the reproductive value (Fisher, 1930) of an individual before dispersal relative to that of an individual after dispersal (i.e. the probability that she survives the dispersal phase) and *r* is the kin selection coefficient of genetic relatedness (Hamilton, 1964) between neighbours before dispersal (see supplementary material for details). Rather than being an independent parameter, relatedness is modulated by demographic details, including rates of voluntary and forced dispersal. In the present model relatedness is given by 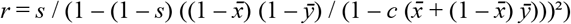, where *s* is the probability that two individuals, randomly chosen from the same patch before dispersal, have the same mother (see supplementary material for details). The degree of sibship *s* reflects factors such as the number of breeding individuals and the degree of reproductive skew within patches. A higher probability of sibship and lower rates of voluntary and/or forced dispersal result in higher relatedness.

Expression (2) is a form of Hamilton’s (1964) rule, and it admits an inclusive-fitness interpretation. Specifically, voluntary dispersal results in: (1) a direct mortality cost *c* paid with probability 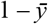 that the actor would not be forced to disperse anyway; (2) a corresponding indirect benefit with the same probability 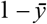, owing to a reduction in the intensity of competition within the actor’s natal patch, where a proportion 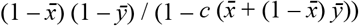 of the beneficiaries are non-dispersing locals who are related to the actor by *r*; (3) an indirect benefit owing to the actor reducing the cost of forcing 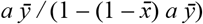 to be paid by her neighbours, who have reproductive value *v* and are related to her by *r*; and (4) an indirect cost owing to corresponding intensification of competition among the actor’s neighbours, who have reproductive value *v* and are related to her by *r*, to the extent 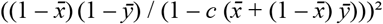 that neighbours compete with each other for reproductive opportunities.

We also find that natural selection favours an increase in forced dispersal if

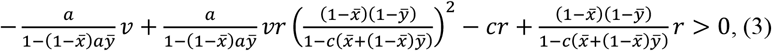

(see supplementary material for details). Forcing a neighbour to disperse leads to: (1) a direct loss of competitiveness for reproductive opportunities 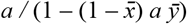, weighted by the actor’s reproductive value *v*; (2) an indirect benefit, owing to a corresponding reduction in the intensity of competition among the actor’s neighbours, who have reproductive value *v* and are related to the actor by *r*, to the extent 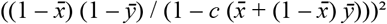 that neighbours compete with each other for reproductive opportunities; (3) an indirect cost of dispersal *c* incurred by the neighbour who has been forced to disperse, and who is related to the actor by *r*; and (4) a corresponding indirect benefit owing to the reduction in the intensity of competition among the actor’s neighbours, who are related to the actor by *r*, to the extent 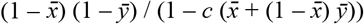 that they remain to compete for reproductive opportunities in the natal patch.

Considering the joint evolution of both voluntary and forced dispersal, we find that— irrespective of the initial state of the population—natural selection always drives the population towards a single stable outcome (see supplementary material for details). Depending on parameter values, this stable outcome may involve (i) voluntary dispersal only, (ii) forced dispersal only, or (iii) a mixture of voluntary and forced dispersal (Figure 1). The “voluntary dispersal only” outcome (*x*\* > 0, *y*\* = 0) obtains when there is a large cost of forcing (high *a*), a low cost of dispersal (low *c*) and a high degree of sibship (high *s*); the “forced dispersal only” outcome (*x*\* = 0, *y*\* > 0) obtains when there is a small cost of forcing (low *a*), a high cost of dispersal (high *c*) and a low degree of sibship (low *s*); and the “mixture of voluntary and forced dispersal” outcome (*x*\* > 0, *y*\* > 0) obtains for a narrow range of intermediate values of the cost of forcing (medium *a*), the cost of dispersal (medium *c*) and the degree of sibship (medium *s*).

**Figure 1.**
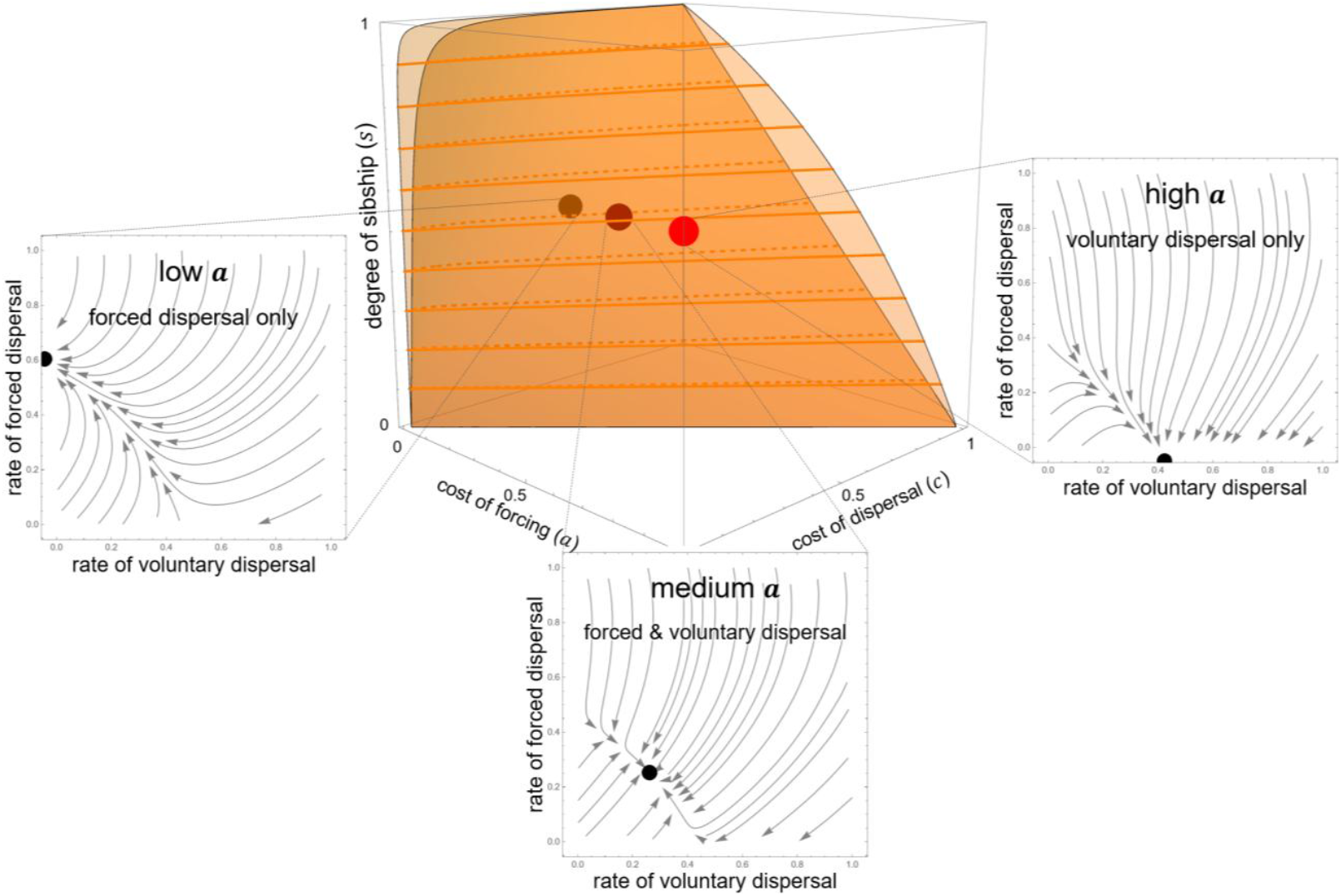
Forms of dispersal, when social modulation does not improve dispersal success. There is only one convergence stable evolutionary outcome for any given model parameterisation (*a, c, s*), represented as a point in space within the 3D plot. For low cost of forcing (low *a*), high cost of dispersal (high *c*) and low degree of sibship (low *s*), only forced dispersal obtains (i.e. *x*\* = 0, *y*\* > 0; portion of parameter space lying behind the orange volume). For high cost of forcing (high *a*), low cost of dispersal (low *c*) and high degree of sibship (high *s*), only voluntary dispersal obtains (i.e. *x*\* > 0, *y*\* = 0; portion of parameter space lying in front of the orange volume). For intermediate cost of forcing (medium *a*), intermediate cost of dispersal (medium *c*) and intermediate degree of sibship (medium *s*), both voluntary and forced dispersal coexist (i.e. *x*\* > 0, *y*\* > 0; portion of parameter space lying within the orange volume). Inset are vector fields illustrating the evolutionary dynamics of voluntary and forced dispersal for each of these three outcomes.

It is intuitive that only voluntary dispersal is favoured when the cost of forcing a neighbour to disperse is relatively high (i.e. high *a*) and the mortality cost of dispersal is relatively low (i.e. low *c*). It is also intuitive that only forced dispersal is favoured when the cost of forcing a neighbour to disperse is relatively low (i.e. low *a*) and the mortality cost of dispersal is relatively high (i.e. high *c*). Less intuitive is that voluntary and forced dispersal can be favoured simultaneously, because if forcing is serving any adaptive purpose then this implies that the resulting level of dispersal is greater than the level the individuals are willing to disperse, which suggests that the level of voluntary dispersal should therefore be favoured to decrease. However, coexistence of voluntary and forced dispersal is possible—across a narrow range of intermediate parameter values—because individuals gain an additional benefit of voluntarily dispersing in the context of forced dispersal as it saves their neighbouring kin from having to pay the cost of forcing their dispersal (see supplementary material for details). Finally, it is intuitive that a population transitions from having only forced dispersal to only voluntary dispersal as the degree of sibship increases (i.e. higher *s*) because, as neighbours become more closely related to each other, they are increasingly favoured to avoid the wasteful cost of forced dispersal. Indeed, in a clonal-patch setting (i.e. *s* = 1) there is no possibility for the evolution of forced dispersal.

In terms of the level of dispersal that is evolutionarily favoured (Figure 2, see supplementary material for details), we find that an increase in the cost of dispersal (higher *c*) is associated with a reduced level of voluntary dispersal (lower *x*\*) and a lower level of dispersal overall (lower *x*\* + (1 – *x*\*) *y*\*)—although it can lead to either an increase or a decrease in the level of forced dispersal (higher or lower *y*\*), depending on parameter values. An increase in the degree of sibship (higher *s*) is associated with an increased level of voluntary dispersal (higher *x*\*) and a higher level of dispersal overall (higher *x*\* + (1 – *x*\*) *y*\*)—though it can lead to either an increase or a decrease in the level of forced dispersal (higher or lower *y*\*), depending on parameter values. Finally, an increase in the cost of forcing a neighbour to disperse (higher *a*) is associated with a decrease in the level of forced dispersal (lower *y*\*) and in the overall level of dispersal (lower *x*\* + (1 – *x*\*) *y*\*)—except within the “voluntary dispersal only” outcome, where it has no effect.

**Figure 2.**
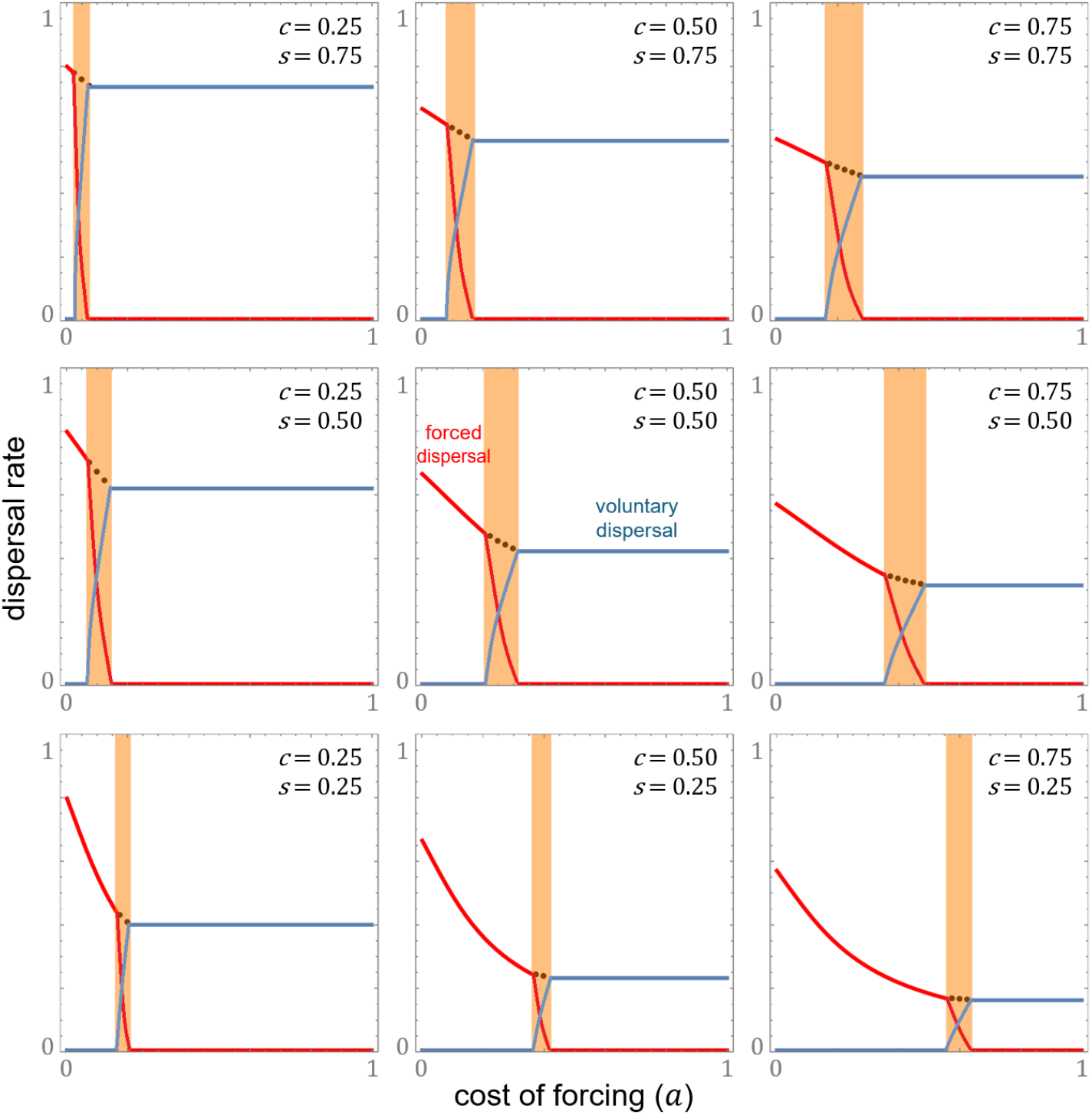
Rates of dispersal, when social modulation does not improve dispersal success. A higher cost of forcing (higher *a*) leads to a decrease in the level of forced dispersal (lower *y*\*, red line) and in the overall level of dispersal (lower *x*\* + (1 – *x*\*) *y*\*). A higher cost of dispersal (higher *c*) leads to a decrease in the level of voluntary dispersal (lower *x*\*, blue line) and a lower level of dispersal overall (lower *x*\* + (1 – *x*\*) *y*\*). A higher degree of sibship (higher *s*) leads to an increase in the level of voluntary dispersal (higher *x*\*) and a higher level of dispersal overall (higher *x*\* + (1 – *x*\*) *y*\*). The black dotted line shows the overall level of dispersal when voluntary and forced dispersal coexist (orange region).

### Facilitated dispersal

Here we use the same lifecycle and population structure as in the first model but assume that the mortality cost incurred by dispersers is lower for socially modulated dispersal than for voluntary dispersal. This may be because the modulation helps the recipient initiate effective dispersal (such as through utilising extrinsic kinetic energy, as in social amoebae) or survive dispersal (such as through protective behaviours and information sharing). In this case, the relative fitness of a focal individual can be written as

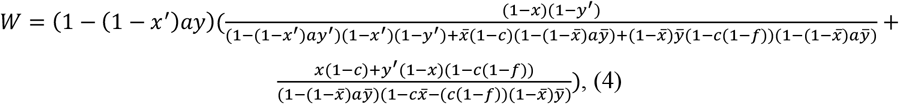

where *f* is the proportion of the mortality cost of dispersal that is alleviated by social modulation (i.e. setting *f =* 0 recovers the model of the previous section).

We find that natural selection favours an increase in voluntary dispersal if

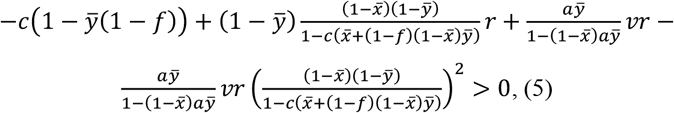

where 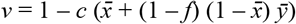 and 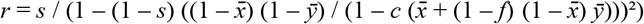 (see supplementary material for details). As a form of Hamilton’s (1964) rule, expression (5) shows that voluntary dispersal results in: (1) a direct mortality cost *c* paid with probability 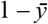 that the actor would not be sent out and incur the reduced dispersal cost *c* (1 – *f*) anyway; (2) a corresponding indirect benefit with the same probability 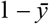, owing to reduced competition within the actor’s natal patch, where a proportion 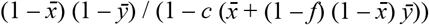 of the beneficiaries are non-dispersing locals who are related to the actor by *r*; (3) an indirect benefit due to the actor reducing the cost of modulation 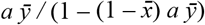 to be paid by her neighbours, who have reproductive value *v* and are related to her by *r*; and (4) an indirect cost owing to corresponding intensification of competition among the actor’s neighbours, who have reproductive value *v* and are related to her by *r*, to the extent 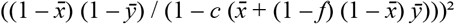 that neighbours compete with each other for reproductive opportunities.

Also, we find that natural selection favours an increase in facilitated dispersal when

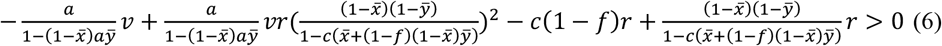

(see supplementary material for details), showing that modulating a neighbour’s dispersal leads to: (1) a direct loss of competitiveness for reproductive opportunities 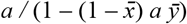, weighted by the actor’s reproductive value *v*; (2) an indirect benefit, owing to corresponding reduced competition among the actor’s neighbours, who have reproductive value *v* and are related to the actor by *r*, to the extent 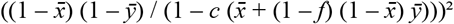 that neighbours compete with each other for reproductive opportunities; (3) an indirect cost of facilitated dispersal *c* (1 – *f*) incurred by the neighbour who has been sent out, who is related to the modulator by *r*; and (4) a corresponding indirect benefit due to the reduction in the intensity of competition within the actor’s natal patch, where a proportion 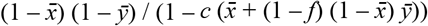 of the beneficiaries are non-dispersing locals who are related to the actor by *r*.

Considering the joint evolution of both forms of dispersal, we find that—as in the model of the previous section—depending on parameter values, there are three stable outcomes: (i) voluntary dispersal only, (ii) facilitated dispersal only, and (iii) a mixture of the two (Figure 3; see supplementary material for details). The “voluntary dispersal only” outcome (*x*\* > 0, *y*\* = 0) obtains when there is a low level of facilitation (low *f*), a large cost of modulation (high *a*), a low cost of dispersal (low *c*) and a high degree of sibship (high *s*); the “facilitated dispersal only” outcome (*x*\* = 0, *y*\* > 0) obtains when there is a high level of facilitation (high *f*), a small cost of modulation (low *a*), a high cost of dispersal (high *c*) and a low degree of sibship (low *s*); and a narrow range of intermediate values of the four parameters may result in the “mixture of voluntary and facilitated dispersal” outcome (*x*\* > 0, *y*\* > 0).

**Figure 3.**
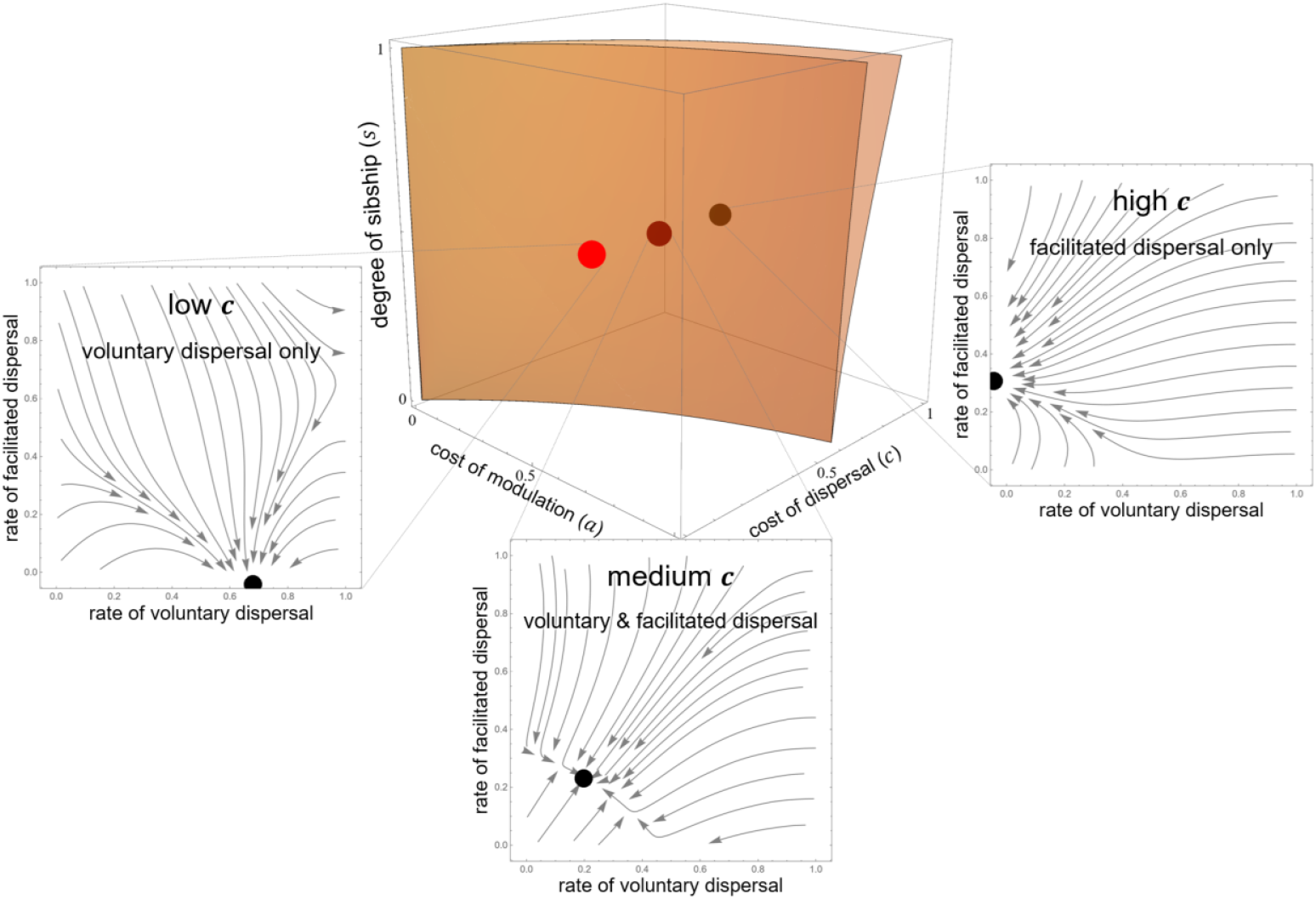
Forms of dispersal, when social modulation improves dispersal success. For low cost of modulation (low *a*), high cost of dispersal (high *c*) and low degree of sibship (low *s*), only facilitated dispersal obtains (i.e. *x*\* = 0, *y*\* > 0; portion of parameter space lying behind the orange volume). For high cost of modulation (high *a*), low cost of dispersal (low *c*) and high degree of sibship (high *s*), only voluntary dispersal obtains (i.e. *x*\* > 0, *y*\* = 0; portion of parameter space lying in front of the orange volume). For intermediate cost of modulation (medium *a*), intermediate cost of dispersal (medium *c*) and intermediate degree of sibship (medium *s*), both voluntary and facilitated dispersal coexist (i.e. *x*\* > 0, *y*\* > 0; portion of parameter space lying within the orange volume). Increasing the level of facilitation (higher *f*) expands the regions where facilitated dispersal is favoured and where voluntary dispersal is disfavoured especially when the cost of dispersal is high (i.e. when the modulation is effective in facilitating dispersal). Here the modulation is set to reduce the dispersal cost by half (*f =* 0.5; see supplementary material for outcomes for different values of *f*).

It is intuitive that only voluntary dispersal is favoured when the cost of modulating a neighbour’s dispersal is high, the level of facilitation that the modulation involves is low and the cost of dispersal is low (i.e. high *a*, low *f &* low *c*) while only facilitated dispersal is favoured when the opposite obtains (i.e. low *a*, high *f &* high *c*). An intermediate region in which both forms of dispersal coexist is, again, possible because in the context of facilitated dispersal voluntary dispersers gain an addition benefit of saving their kin from having to pay the cost of modulating their dispersal. As in the previous model, as the degree of sibship increases (i.e. higher *s*) individuals are more likely to exhibit only voluntary dispersal because they are increasingly favoured to avoid the cost of modulation. However, because facilitated dispersal lowers the cost of dispersal incurred by the recipient and thus have an extra indirect benefit, it can now be favoured even in a clonal-patch setting (*s =* 1) if it is sufficiently effective (e.g. low *a*, high *f &* high *c*). Additionally, because modulation not only benefits the non-dispersing kin of the actor but also the disperser in terms of a reduced mortality cost, facilitated dispersal can be favoured even if the cost of modulation is higher than the cost of dispersal itself (Figure 3; see supplementary material for details).

In terms of the level of dispersal that is evolutionarily favoured (Figure 4; see supplementary material for details), we find that—as in the model of the previous section—an increase in the cost of dispersal (higher *c*) and a decrease in the degree of sibship (lower *s*) both lead to a lower level of voluntary dispersal (lower *x*\*) and a lower level of dispersal overall (lower *x*\* + (1 – *x*\*) *y*\*)—though they can lead to either an increases or a decrease in the level of facilitated dispersal (higher or lower *y*\*), depending on parameter values. What is different is that, although a decreased cost of modulating a neighbour’s dispersal (lower *a*) is associated with an increased level of facilitated dispersal (higher *y*\*), it can lead to either an increase or a decrease in the overall level of dispersal (higher or lower *x*\* + (1 – *x*\*) *y*\*), depending on parameter values. This is because, while the parameter region favouring modulated dispersal expands as this facilitation benefit increases, this facilitation also increases the success rate of dispersal, which weakens selection for dispersal by lowering stabilised relatedness among neighbours and the share of reduced competition that the non-dispersing neighbours enjoy. Therefore, with intermediate parameter values (e.g. intermediate *a*), adoption of dispersal facilitation can, counterintuitively, be associated with a reduced overall level of dispersal. For the same reason, while an increase in the level of facilitation (higher *f*) leads to a higher level of facilitated dispersal (higher *y*\*) it can lead to an increase or a decrease in the overall level of dispersal (higher or lower *x*\* + (1 – *x*\*) *y*\*).

**Figure 4.**
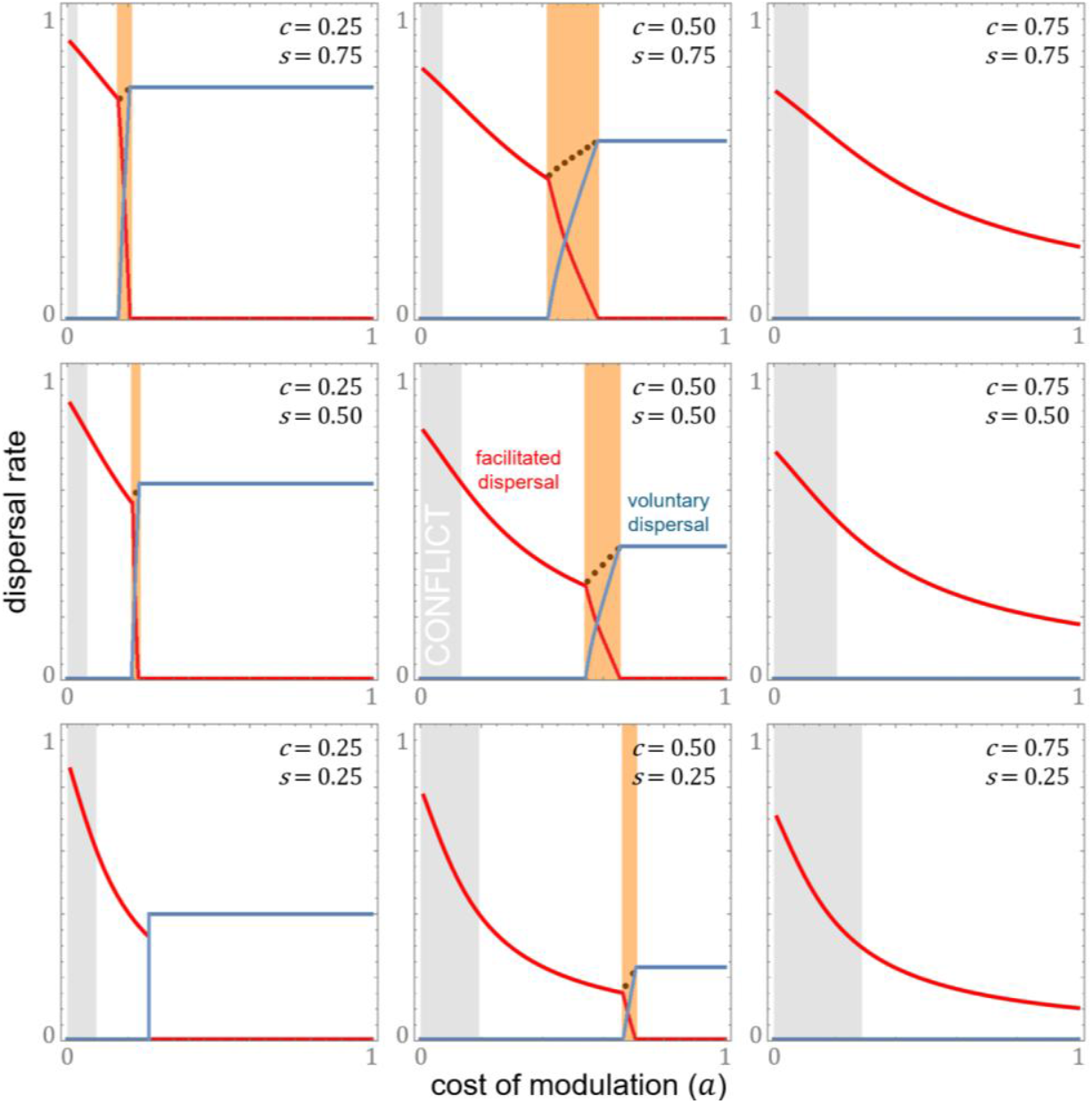
Rates of dispersal, when social modulation improves dispersal success. A higher cost of dispersal (higher *c*) and a lower degree of sibship (lower *s*) both lead to a decrease in the level of voluntary dispersal (lower *x*\*, blue line) and a lower level of dispersal overall (lower *x*\* + (1 – *x*\*) *y*\*). Both a lower cost of modulation (lower *a*) and a higher level of facilitation (higher *f*) lead to an increase in the level of facilitated dispersal (higher *y*\*, red line), but they can lead to an increase or a decrease in the level of dispersal overall (higher or lower *x*\* + (1 – *x*\*) *y*\*). Because it reduces the cost incurred by dispersers, the modulation may be welcomed (non-grey region) or opposed (grey region) by the recipient. The black dotted line shows the overall level of dispersal when voluntary and facilitated dispersal coexist (orange region). For illustration, we assume that modulation halves the mortality cost of dispersal (*f =* 0.5).

In contrast to the model of the previous section, in which modulated dispersal is always counter to the inclusive-fitness interests of the recipient (i.e. “forced” dispersal), we find that facilitated dispersal can sometimes be in line with the recipient’s interests (see supplementary material for details). Because individuals can prefer being sent out and paying a lower dispersal cost over voluntarily dispersing and incurring a larger cost, whether there is a conflict between the actor and the recipient of the modulation depends on the level of modulated dispersal. Only when the cost of modulation is so low that modulated dispersal occurs at a higher rate than that they are favoured to disperse at the reduced cost, there will be an evolutionary conflict of interest. For instance, when the cost of modulating another’s dispersal is larger than the cost incurred by the modulated disperser (i.e. *a* > *c* (1 – *f*)), modulation will be welcomed by the recipients. Only in the non-conflict region, the adoption of modulated dispersal can lead to a decreased overall rate of dispersal.

## Discussion

Dispersal across the biological world need not be under the sole control of the disperser, but may in some scenarios also be modulated by her social partners. Employing kin selection methodology, we have performed a mathematical analysis of the conditions under which individuals are favoured to exert an influence on their social partners’ dispersal and how the evolution of such modulated dispersal affects the evolution of voluntary dispersal as well as the level of dispersal overall. We have shown that: (1) there are three possible stable evolutionary outcomes, involving only voluntary dispersal, only modulated dispersal, or both forms of dispersal; (2) these outcomes depend on the interplay between the values of four model parameters, describing the cost of dispersal, the cost of modulation, the level of facilitation, and the degree of sibship; (3) facilitated dispersal may, counterintuitively, lead to a reduction in the overall level of dispersal; and (4) social modulation may be welcome or unwelcome from the perspective of the disperser.

We have shown that if individuals are able to modulate the dispersal of their social partners they may be favoured to do so and, if this is the case, then it may either completely replace or else coexist with voluntary dispersal. Hamilton & May’s (1977) analysis of purely voluntary dispersal demonstrates that this can be favoured by natural selection even if it reduces the direct fitness of the disperser because it tends to reduce the local competition for her kin (see also Frank, 2013). Here we show that voluntary dispersal will only be favoured when the cost of modulation is relatively high. This is because the favoured rate of voluntary dispersal is usually lower than the group optimum, which incentivises individuals to exert an influence on each other’s dispersal. We have found that, in the context of modulated dispersal, voluntary dispersal yields an extra indirect fitness benefit because it saves the individual’s neighbouring kin from having to pay the cost of modulating her dispersal, and this enables the two forms of dispersal to coexist within an intermediate region of parameter space.

Considering the joint evolution of voluntary and socially modulated dispersal, we have found that, ultimately, only modulated dispersal obtains when there is a high mortality cost of dispersal, a low degree of sibship among neighbours, a low cost of modulating dispersal and that modulation substantially lowers the cost of dispersal—and we find that only voluntary dispersal obtains under the opposite condition. In particular, if the social modulation does not facilitate dispersal (i.e. does not reduce the mortality cost of dispersal), then it cannot be favoured when the cost of modulation is greater than the mortality cost of dispersal itself or when group mates are all clonally related (because in this case the group-optimal level of dispersal is attained on a purely voluntary basis). Yet, if modulation does substantially facilitate dispersal, then it can be favoured even in a clonal-group setting and even when it is so costly as to be effectively fatal to the modulator. One classic example of socially modulated dispersal is that of social amoebae, in which some cells make the ultimate sacrifice to form a stalk that facilitates the dispersal of their social partners as spore when starvation is imminent (Bonner, 2015). Here, we have clarified that neither clonality nor heterogeneity in patch quality is essential to the evolution of facilitated dispersal.

Concerning modulation that does not facilitate dispersal—i.e. does not reduce the mortality cost of dispersal—we find that the adoption of modulated dispersal results in a higher level of dispersal overall. Yet, if modulation facilitates dispersal, its adoption may increase or decrease the overall level of dispersal. This is because a reduction in the mortality cost of dispersal increases the proportion of dispersing individuals that successfully reach their destinations, which reduces the intensity of kin competition which is ultimately the driver of dispersal. Hence, when the cost of modulation is low the adoption of facilitated dispersal increases the overall level of dispersal, but when the cost of modulation is intermediate it may reduce the overall level of dispersal. In line with previous analyses of dispersal in stable habitats (Frank, 1986; Hamilton & May, 1977; Taylor, 1988), we find that an increase in the cost of dispersal and a decrease in relatedness among neighbours both lead to a lower level of voluntary dispersal and a lower level of dispersal overall.

Finally, we have found that modulated dispersal—depending on whether and the extent to which it reduces the mortality cost of dispersal—may or may not trigger an inclusive-fitness conflict between the actor and the recipient. Hypothetically granting the recipients control over whether or not they do disperse in response to modulation (see supplementary material for details), we find that a higher level of facilitation increases recipients’ willingness to be dispersed as they thereby reduce local competition for their kin at a lower cost than they can achieve with voluntary dispersal. Accordingly, while there is conflict when the level of facilitation is low, the interaction may be cooperative when the level of facilitation is higher. This finding suggests that one possible way a parent can manipulate the dispersal tendencies of their offspring is by equipping them with a phenotype that has better dispersal success (cf. Rodrigues & Gardner, 2022).

This distinction between welcomed and unwelcomed modulation may be crucial for understanding the evolution of dispersal, as the capacity to force and to facilitate may be very different for the same species, which would predispose some taxa to one form of modulated dispersal rather than the other. For instance, a dominant chimpanzee might easily evict a subordinate but might not be able to effectively improve how successfully it disperses, whereas a group of microbes that are unable to exclude any cells might be able to aggregate and assist in some of the cells’ passive dispersal (Bonner, 2015; Kaiser, 2004). Also, if the modulation acts against the recipient’s inclusive-fitness interests, counteradaptation may arise, which can lead to a coevolution of manipulation and resistance, whereas if the modulation is in line with the recipient’s inclusive-fitness interests, it might readily develop into a complex and coordinated social phenotype—as in the cooperative dispersal of social amoebae and other microbes (Bonner, 2015; Engelhardt et al., 2022; Kaiser, 2004; Xue et al., 2011).

Socially modulated dispersal has previously received some attention, particularly in the context of parent-offspring conflict over offspring dispersal. Following Hamilton & May’s (1977) analysis of kin-selected dispersal, Motro (1982a; 1982b; 1983) showed that if parents control their offspring’s dispersal then this is often expected to evolve to a different level than would be the case when dispersal is under the control of the offspring themselves (see also Frank, 2013; Starrfelt & Kokko, 2010). Rodrigues & Gardner (2016) have extended this principle to scenarios involving alloparental control of dispersal. In contrast to those previous analyses which focused on cross-generational modulation of dispersal, we have shown that conflicts over dispersal also exist between same-generation social partners, on account of a basic public-goods dilemma in which all individuals agree that some level of dispersal is good for the group—but all would prefer that someone else does the dispersal. Moreover, we have considered the impact of the cost of modulation and associated benefits for dispersers, going beyond the battleground analysis (cf. Godfray, 1995) that diagnoses conflicts of interest to consider the coevolutionary dynamics of modulated and voluntary dispersal to determine how these conflicts are ultimately resolved (cf. Godfray, 1995).

As an initial exploration of the widely observed phenomenon of socially modulated dispersal, we have aimed for a generic and illustrative analysis. Firstly, for the sake of facilitating synthesis of new results with the existing literature, we have adopted Hamilton & May’s (1977) framework and inherited several of their basic model assumptions. We have made the usual assumption that the number of pre-dispersal juveniles is large (Frank, 1986; Hamilton & May, 1977; Taylor, 1988; 1992), and one consequence of this is that when an individual causes a neighbour to disperse this does not meaningfully reduce the local competition experienced by the modulator but rather this benefit accrues to her neighbours. If modulators stood to gain direct benefits then this would be expected to promote the evolution of modulated dispersal. Also, we have assumed the infinite-island structure of Wright (1931), in which successful dispersers choose destinations randomly from an effectively infinite number of patches and thus always disperse into a patch without relatives. Relaxing this assumption should weaken the kin-competition alleviating effect of dispersal and thus reduce selection for both voluntary and socially modulated dispersal, but it is not expected to alter the qualitative findings of the current analysis. These assumptions are less applicable to species which exhibit low fecundity and are particularly selective in choosing destination patch; future research that relaxes these assumptions may better capture the biology of those systems.

Relatedly, incorporating socially modulated dispersal into the classic framework, we have assumed that—regardless of the form of dispersal—successful dispersers individually relocate to pre-existing patches, which offer essentially the same reproductive opportunities they would have enjoyed had they not dispersed. Although individual dispersal has been shown to be particularly favoured for competition avoidance (Soubeyrand et al., 2015), dispersal facilitation might not only reduce the cost of dispersal but could also help dispersers discover or invade unoccupied territories and dispersers might relocate collectively to shared destinations as a result of facilitation (Maag et al., 2022; Ridley, 2011). Collective dispersal has been proposed as an evolutionary pathway for the evolution of altruism as it may preserve the close relatedness between social partners (Gardner & West, 2006; Rodrigues & Taylor, 2018). Future work could investigate the evolution of modulated dispersal in the context of collective dispersal and how it may influence the group dynamics and potential for altruism within a population.

Lastly, we have assumed that modulation of dispersal operates in an indiscriminate manner, which relies on no complex social structure or kin discrimination. In a heterogeneous population, however, peer-to-peer modulation could be practised in a way that is based upon neighbours’ characteristics (e.g. sex, cooperativeness or competitiveness) or their kinship or other social affiliation to the modulator. In such cases, modulation might greatly alter demography and population dynamics. For example, when mating competition is stronger in one sex, modulated dispersal may be favoured to operate in a sexually biased way. Thus, a tendency for the less-dispersing sex to give and receive more altruism might arise, as they are more related to the group pro-dispersal (Micheletti et al., 2020). Equally, in a heterogeneous population, different individuals (e.g. depending on dominance or age) may be favoured to exhibit distinct modulation strategies (Sarno et al, 2003; Strickland, 1991; Thompson et al., 2017). Future exploration that extends our analysis and examines the knock-on effects of the emergence of modulated dispersal will help advance our understanding of the social evolution of dispersal.

