## supplementarymaterial for "The evolution of socially modulated dispersal"

Forced dispersal

*Fitness function*—We assume an inelastic population structure, in which the number of juveniles at the beginning of a lifecycle remains unchanged across generations. Hence, the expected number of offspring that reach the beginning of the next cycle is one per individual (*w̅* = 1). We can then express the relative fitness of the focal individual in terms of how many offspring she is expected to have (*W* = *w*/*w̅* = *w*/1 = *w*).

Under the assumptions of homogeneity in patch quality and that dispersers choose destinations randomly, we can write the relative fitness as

$W=(1-\left( 1-x^{'} \right)ay)(\left( 1-x \right)\left( 1-y^{'} \right)\frac{n}{\left( 1-\left( 1-x^{'} \right)ay^{'} \right)\left( 1-x^{'} \right)\left( 1-y^{'} \right)n+\left( 1-c \right)\left( 1-\left( 1-\bar{x} \right)a\bar{y} \right)\left( \bar{x}+\left( 1-\bar{x} \right)\bar{y} \right)n}+\left( 1-c \right)\left( x+\left( 1-x \right)y^{'} \right)\frac{n}{\left( 1-\left( 1-\bar{x} \right)a\bar{y} \right)\left( 1-c\left( \bar{x}+(1-\bar{x} \right)\bar{y} \right))n})$. (S1)

The first set of outermost brackets is the proportion of competitiveness for reproductive opportunities remaining after the expected reduction due to forcing and the rest (i.e. the second set of outermost brackets) is the expected number of her offspring before the adjustment. The first set of brackets is subject to the local voluntary dispersal rate because the number of nondispersers determines the number of potential recipients of forcing; for instance, if *x' =* 1 (i.e. full voluntary dispersal) no modulation will be possible, and, if *x' =* 0.5 (half of the group are dispersing) there will be only one potential recipient in every two individuals. The level of modulation (*y,* *y', ȳ*) can be interpreted as either a probabilistic tendency of modulating or a degree of participation in a shared effort.

For the second set of outermost brackets, if the focal individual does not voluntarily disperse (with probability 1 – *x*) and has not been forced to disperse (with probability 1 – *y'*, see eqn S2), she will share all reproductive opportunities (i.e. *n*, the carrying capacity of a patch; see the first fraction) with (1) her original neighbours (i.e. patch mates) that managed to stay and, potentially, lost some competitiveness due to forcing and (2) successful dispersers, who dispersed into the group either due to their voluntary or involuntary dispersal and may have lost some competitiveness due to their forcing behaviour in their own natal patch. Otherwise, if the focal individual has not died during dispersal (with probability 1 – *c*), she will share *n* with the nondispersers there and the other newcomers in that patch (i.e. the second fraction). Simplifying eqn S1 results in eqn 1 in the main text.

The number of times that modulation happens in a patch equals the average level of forcing implemented within the group multiplied by the number of individuals in the patch (i.e. $n$). Dividing that by the number of individuals that do not voluntarily disperse gives the rate of forced dispersal in the focal patch:

$\frac{(\left( 1-x^{'} \right)y^{'})n}{(1-x^{'})n}=y^{'}$, (S2)

except for full voluntary dispersal (i.e. where forcing is impossible).

*Hamilton’s rule & relatedness*—Considering a locus that controls the voluntary dispersing behaviour, we may write the marginal fitness of voluntary dispersal as

$\frac{dW}{dg}=\frac{\partial W}{\partial x}\frac{dx}{dG}\frac{dG}{dg}+\frac{\partial W}{\partial x^{'}}\frac{dx^{'}}{dG^{'}}\frac{dG^{'}}{dg}$, (S3)

where $g$ is the genic value of a gene drawn from this locus in the focal individual, *G* is the average genic value of the genes from the locus for the focal individual and *G'* is that for her neighbours. *dG/d*$g$ is the consanguinity of the focal individual to herself while *dG'/d*$g$ is the consanguinity of the focal individual to the neighbours, and the mapping between genotype and phenotype is *dx/dG = dx'/dG'* (Taylor & Frank, 1996; see also Gardner et al., 2011). That is, if eqn S3 is positive (*dW/d*$g$ *>* 0) voluntary dispersal is favoured by natural selection to increase (and if eqn S3 is negative it is disfavoured). From here, we may simplify this inequality by dividing both sides by the mapping and the consanguinity of the focal individual to herself, which gives

$\frac{\partial W}{\partial x}+\frac{\partial W}{\partial x^{'}}r>0$, (S4)

where *r* is the kin-selection coefficient of relatedness between neighbours.

Under the assumption of vanishingly low variation in traits values in the population, the condition for an increase in voluntary dispersal is then

$-\frac{c\left( 1-\bar{y} \right)}{1-c\left( \bar{x}+\left( 1-\bar{x} \right)\bar{y} \right)}+\frac{\left( 1-\bar{x} \right)\left( 1-\bar{y} \right)^{2}}{\left( 1-c\left( \bar{x}+\left( 1-\bar{x} \right)\bar{y} \right) \right)^{2}}r+\frac{a\bar{y}}{\left( 1-\left( 1-\bar{x} \right)a\bar{y} \right)}r-\frac{a{\bar{y}\left( 1-\bar{x} \right)}^{2}\left( 1-\bar{y} \right)^{2}}{\left( 1-\left( 1-\bar{x} \right)a\bar{y} \right)\left( 1-c\left( \bar{x}+\left( 1-\bar{x} \right)\bar{y} \right) \right)^{2}}r>0$, (S5)

which is rearranged into Expression 2 in the main text for conceptual clarity—by multiplying both sides the reproductive value of an individual before dispersal relative to that of an individual after dispersal (i.e. 1 – *c*(*x̄ +* (1 – *x̄*)*ȳ*); *v* in the main text).

Using the same method, we obtain the condition for forced dispersal to increase:

$-\frac{a(1-\bar{x})}{1-(1-\bar{x})a\bar{y}}+\frac{a{(1-\bar{x})}^{3}{(1-\bar{y})}^{2}}{(1-(1-\bar{x})a\bar{y}){(1-c(\bar{x}+(1-\bar{x})\bar{y}))}^{2}}r-\frac{c(1-\bar{x})}{1-c(\bar{x}+(1-\bar{x})\bar{y})}r+\frac{{(1-\bar{x})}^{2}(1-\bar{y})}{{(1-c(\bar{x}+(1-\bar{x})\bar{y}))}^{2}}r>0$, (S6)

which is rearranged into Expression 3 in the main text.

Setting *ȳ* = 0, Expression S5 recovers results from existing models of kin-selected dispersal (Frank, 1986; Hamilton & May, 1977) including that voluntary dispersal is favoured when relatedness between neighbours is higher than the cost of dispersal (*r > c*$)$ and, in particular, stabilises at

$x^{*}=\left\{ \begin{aligned} 0 &if r\leq c \\ \frac{r-c}{r-c^{2}} &if r>c \end{aligned} \right.$. (S7)

Rather than being an independent parameter, relatedness is subject to rates of voluntary and forced dispersal. In this model, this is given by

$r=s+(1-s){(\frac{(1-\bar{x})(1-\bar{y})}{1-c(\bar{x}+(1-\bar{x})\bar{y})})}^{2}r$, (S8)

where *s* is the probability that two individuals, randomly chosen from the same patch before dispersal, are siblings. Solving eqn S8 for *r* gives the relatedness listed in the main text (i.e. *r* = *s* / (1 – (1 – *s*) ((1 – *x̄*) (1 – *ȳ*) / (1 – *c* (*x̄* + (1 – *f*) (1 – *x̄*) *ȳ*)))²) ). Setting *x̄* = 0 & *ȳ* = 0 gives *r* = 1. So, unless dispersal is literally not achievable (i.e. *c* = 1), zero dispersal is not a stable outcome.

*Rates & conditions*—Letting relatedness change according to dispersal and setting *ȳ* = 0 in Expression S5, we obtain the marginal fitness of voluntary dispersal when it is the only form of dispersal favoured. Setting the left-hand side of this substituted Expression S5 to zero and solving for *x̄*, we obtain the stabilised rate of voluntary dispersal when it is the only form favoured:

$x^{*}=\frac{s}{c+\frac{s}{2}(1+\sqrt{1+\frac{4c^{2}(1-s)}{s^{2}}})}$, (S9)

which increases with a higher degree of sibship *s* and a lower cost of dispersal *c*. Setting *s* = 1, we recover the group optimum of dispersal in the inelastic stable population structure 1 / (1 + *c*) (Hamilton & May, 1977).

Substituting this rate S9 of voluntary dispersal and its corresponding stabilised relatedness into Expression S6 and setting *ȳ* = 0, we find the condition that forced dispersal is disfavoured to emerge and thus “voluntary dispersal only” is a stable outcome:

$a>\frac{c(2-s(1+\sqrt{1+\frac{4c^{2}(1-s)}{s^{2}}}))}{2(1-c)}$, (S10)

which is always fulfilled when the degree of sibship is one (*s =* 1) and when the cost of modulation is not lower than the cost of dispersal (*a* ≥ *c*). When *s <* 1 & *a* < *c*, the right-hand side of the inequality is always positive, so that forcing will be favoured if its cost is low enough. The right-hand side increases with a higher cost of dispersal and a lower degree of sibship.

Then, letting relatedness change according to dispersal and setting *x̄* = 0 in Expression S6, we find the marginal fitness of forced dispersal when it is the only form of dispersal favoured. Setting the left-hand side of this substituted Expression S6 to zero and solving for *ȳ*, we find the stabilised rate of forced dispersal when it is the only form favoured:

$y^{*}=\frac{2s}{2a+(1+a+c)s+\sqrt{4a^{2}(1+s)+{(1-a+c)}^{2}s^{2}}}$, (S11)

which increases with a higher degree of sibship, a lower cost of dispersal and a lower cost of forcing. In the limit of costless forcing (*a* $≐$ 0) forced dispersal will stabilise at the group optimum 1 / (1 + *c*) if there is no voluntary dispersal, and no parameter changes can result in a higher rate. This shows that, the reason why forced dispersal is not favoured in a clonal-patch setting is that, the individuals are already favoured to achieve the group optimum on a voluntary basis.

Substituting this rate S11 of forced dispersal and its corresponding stabilised relatedness into Expression S5 and setting *x̄* = 0, we find the threshold value of the cost of forcing for voluntary dispersal to be favoured to emerge *a* > *f* (*c, s*), which increases with a lower degree of sibship and a higher cost of dispersal. This condition is more stringent than Expression S10, so that it is always fulfilled when *s =* 1 and *a* ≥ *c*. Yet, similar to Expression S10, when *s <* 1 a low enough cost of forcing can always fulfil the condition. Also, we find that in the region where only forced dispersal is favoured *a* ≤ *f* (*c, s*) *y** is always higher than the would-be rate of voluntary dispersal if forcing is nonexistent (i.e. eqn S9; see Figure S1) while in the region where only voluntary dispersal is favoured (i.e. Expression S10) *x** is always higher than the would-be rate of forced dispersal if voluntary dispersal is nonexistent (i.e. eqn S11).

Letting relatedness change according to dispersal in Expression S6 and leaving *x̄* unspecified, we then obtain the marginal fitness of forced dispersal that applies to the intermediate parameter space where both forms of dispersal coexist. Setting the left-hand side of this substituted Expression S6 to zero and solving for *ȳ*, we obtain the stabilised rate of forced dispersal as a function of the rate of voluntary dispersal. Letting relatedness change according to dispersal and substituting that rate of forced dispersal into Expression S5, we obtain the marginal fitness of voluntary dispersal. Setting the left-hand side of this substituted Expression S5 to zero and solving it for *x̄* gives the stabilised rate of voluntary dispersal *x* = F*(a, c, s), which is a function of the cost of forcing, the cost of dispersal and the degree of sibship and increases with a higher degree of sibship and a lower cost of dispersal. Substituting that rate of voluntary dispersal into the rate of forced dispersal that depends on the rate of voluntary dispersal gives the rate that is a function of the cost of forcing, the cost of dispersal and the degree of sibship *y* = F*(a, c, s), which increases with a lower cost of forcing. Finally, adding the rate of voluntary dispersal and the rate of forced dispersal multiplied by the proportion of individuals that do not voluntarily disperse gives the overall level of dispersal (i.e. *x* +* (1 – *x**) *y**) in the coexistence region.

Finally, knowing that the rate of forced dispersal when it is the only form of dispersal (eqn S11) also increases with a higher degree of sibship and a lower cost of dispersal and that the rate of voluntary dispersal when it is the only form of dispersal (eqn S9) is always higher than the would-be rate of forced dispersal, we know that the adoption of forced dispersal will not change the monotonic relationships between the overall rate of dispersal and (1) dispersal cost and (2) sibship. That is, the overall dispersal rate increases with a higher degree of sibship and a lower cost of dispersal. Additionally, knowing that the rate of forced dispersal when it is the only form of dispersal increases with a lower cost of forcing and this rate is always higher than the would-be rate of voluntary dispersal in that region, we know that the overall dispersal rate increases with a lower cost of forcing (except for the region where only voluntary dispersal is favoured).

*Nondispersers modulating each other*—The analysis above assumes that voluntary and forced dispersal are two independent traits and allows all individuals to force their neighbours. Here, we model the evolution of forced dispersal in a way that only individuals that are not voluntarily dispersing get to force other nondispersers to disperse and see whether the change leads to qualitative differences. This modelling difference could be conceptually understood as forcing taking place after voluntary dispersal or that the voluntary-disperser phenotype is not favoured to perform any forcing.

Under the same set of assumptions, we may write the relative fitness of a focal individual as

$W=\frac{(1-x)(1-y^{'})(1-ay)}{(1-x^{'})(1-y^{'})(1-ay^{'})+(1-c)(\bar{x}+\left( 1-\bar{x} \right)\bar{y}\left( 1-a\bar{y} \right))}+\frac{x\left( 1-c \right)+(1-x)y^{'}\left( 1-ay \right)\left( 1-c \right)}{\left( 1-\bar{x} \right)\left( 1-\bar{y} \right)\left( 1-a\bar{y} \right)+(1-c)(\bar{x}+\left( 1-\bar{x} \right)\bar{y}\left( 1-a\bar{y} \right))}$. (S12)

Here, the loss of competitiveness for reproductive opportunities is *ay* instead of (1 – *x'*) *ay*, given that only individuals that do not voluntarily disperse can force other nondispersers now. Also, if the focal individual is dispersing voluntarily (with probability *x*; see the second fraction), she will be unable to force and thus keeping her full competitiveness.

Using the kin-selection methodology of Taylor & Frank (1996), we find that an increase in voluntary dispersal is favoured when

$-c(1-\bar{y})+a\bar{y}\left( 1-c\bar{y} \right)+\frac{(1-\bar{x}){(1-\bar{y})}^{2}{(1-a\bar{y})}^{2}}{(1-\bar{x})(1-\bar{y})(1-a\bar{y})+(1-c)(\bar{x}+(1-\bar{x})\bar{y}(1-a\bar{y}))}r>0$, (S13)

which the first term is the direct cost of dispersal paid with probability 1 – *ȳ* that the actor would not be forced to disperse anyway, the second term is a direct benefit owing to the cost of forcing *aȳ* saved multiplied by the probability that that cost would not have been neutralised (i.e. one minus the probability that she would get forced to disperse and die if she did not voluntary disperse), and the third term is an indirect benefit earned by probability 1 – *ȳ* that the actor would not be forced to disperse anyway, multiplied by the proportion of competitiveness left after her expected forcing behaviour if she stayed 1 – *aȳ*—owing to reduced competition within her natal patch, where proportion (1 – *x̄*) (1 – *ȳ*) of her neighbours are expected to stay and have lost some competitiveness due to their forcing behaviour and are competing with themselves, newcomers that succeeded forced dispersal and lost some competitiveness in their natal patch and newcomers that voluntarily dispersed and have not lost any competitiveness on forcing. Again, setting *ȳ =* 0, it recovers the classic findings on kin-selected dispersal (Frank, 1986; Hamilton & May, 1977):

$x^{*}=\left\{ \begin{aligned} 0, &r\leq c \\ \frac{r-c}{r-c^{2}}. &r>c \end{aligned} \right.$ (S14)

In this model, the relatedness between neighbours is given *r* = *s +* (1 – *s*) ((1 – *x̄*) (1 – *ȳ*) (1 – *aȳ*) / ((1 – *aȳ*) (1 – *x̄*) (1 – *ȳ*) + (1 – *c*) ((1 – *x̄*) *ȳ* (1 – *aȳ*) + *x̄*)))² *r*. Solving this for *r* gives

$r=\frac{s}{1-{(1-s)(\frac{(1-\bar{x})(1-\bar{y})(1-a\bar{y})}{(1-\bar{x})(1-\bar{y})(1-a\bar{y})+(1-c)(\bar{x}+(1-\bar{x})\bar{y}(1-a\bar{y}))})}^{2}}$, (S15)

which gives *r =* 1 when *x̄* = 0 & *ȳ =* 0 and thus shows that zero dispersal is an unstable outcome.

Then, following the steps in the previous section, we find the stabilised rate of voluntary dispersal when it is the only form of dispersal favoured and its corresponding parameter space, which are both identical to the results of the previous model (i.e. eqn S9 and Expression S10). So that, forced dispersal can only be favoured when *s <* 1 & *a < c*, and when *s <* 1 forcing will be favoured when the cost of forcing is sufficiently low.

In this model, the condition for natural selection to favour an increase in forced dispersal is

$-a\left( 1-c\bar{y} \right)+a\left( 1-\bar{y} \right)\frac{\left( 1-\bar{x} \right)\left( 1-\bar{y} \right)\left( 1-a\bar{y} \right)}{(1-\bar{x})(1-\bar{y})(1-a\bar{y})+(1-c)(\bar{x}+(1-\bar{x})\bar{y}(1-a\bar{y}))}r-c\left( 1-a\bar{y} \right)r+\left( 1-a\bar{y} \right)\frac{\left( 1-\bar{x} \right)\left( 1-\bar{y} \right)\left( 1-a\bar{y} \right)}{(1-\bar{x})(1-\bar{y})(1-a\bar{y})+(1-c)(\bar{x}+(1-\bar{x})\bar{y}(1-a\bar{y}))}r>0$, (S16)

which the first term is a direct loss of competitiveness for reproductive opportunities if the focal individual has not been forced to disperse herself and died, the second term is an indirect benefit if the actor has not been forced to disperse, owing to corresponding reduced competition in her natal patch, where proportion (1 – *x̄*) (1 – *ȳ*) of her neighbours are expected to stay and have lost some competitiveness due to their forcing behaviour and are competing with themselves, newcomers that succeeded forced dispersal and lost some competitiveness in their natal patch and newcomers that voluntarily dispersed and have not lost any competitiveness on forcing, the third term is an indirect cost of dispersal incurred by the recipient of the forcing, who as an individual who has not voluntarily dispersed lost some competitiveness due to her forcing behaviour, and the fourth term is an indirect benefit multiplied by the remaining competitiveness of the recipient 1 – *aȳ*—owing to corresponding reduced competition for her neighbours.

Then, following the steps in the previous section, we find the same stabilised rate of forced dispersal when it is the only form of dispersal favoured (i.e. eqn S11) and the corresponding parameter space, which is slightly smaller than the one in the first model and is a subregion of the parameter space that violates Expression S10. This threshold value of *a* for forced dispersal to be the only form of dispersal, again, increases with a lower degree of sibship and a higher dispersal cost and is always positive when *s <* 1. Hence, this difference does not change the qualitative findings that (1) in the context of forced dispersal voluntary dispersal earns an extra benefit owing to a reduced cost of forcing to be paid and thus there is an intermediate region between the conditions where only one form of dispersal is favoured, and, on (2) the monotonic relationships between the overall dispersal rate and the three model parameters *a, c* & *s*.

*Sexual reproduction*—Here we investigate the evolution of socially modulated dispersal under haploidy and diploidy in a population that exhibits sexual reproduction.

Specifically, we assume an infinite, patch-structured population where each patch contains a large number of males and females, who mate at random within their patch. Each female mates once while each male mates potentially numerous times. Males then die, and each female either undertakes voluntary dispersal or does not. Some females that do not disperse voluntarily may be forced to disperse by their neighbours. If a disperser survives dispersal, she reaches a randomly chosen patch. Following dispersal, both original inhabitants and surviving incoming dispersers compete for reproductive opportunities within their patch; those that are successful produce a large number of offspring, and then all adults die. This returns the population to the beginning of the lifecycle.

Considering a locus that controls the voluntary dispersing behaviour, we may write the marginal fitness of voluntary dispersal as

$\frac{dW}{dg}=c_{f}\frac{dW_{f}}{dg_{f}}+c_{m}\frac{dW_{m}}{dg_{m}}$, (S17)

where $c_{f}$ and $c_{m}$ are the class reproductive values (Fisher, 1930) for females and males, respectively, and ${dW}_{f}/dg_{f}$ and ${dW}_{m}/dg_{m}$ are the marginal fitness of voluntary dispersal in females and males. The class reproductive values are equal between the two sexes here (i.e. $c_{f}$ = $c_{m}$) in both haploidy and diploidy, so we may simplify the direction of natural selection acting upon voluntary dispersal as

$\frac{dW_{f}}{dg_{f}}+\frac{dW_{m}}{dg_{m}}$, (S18)

which can be expanded into

$(\frac{\partial W_{f}}{\partial x}p_{f}+\frac{\partial W_{f}}{\partial x^{'}}p_{ff})+(\frac{\partial W_{m}}{\partial x}p_{m}+\frac{\partial W_{m}}{\partial x^{'}}p_{mf})$, (S19)

where $p_{f}$ is the consanguinity to self from a female perspective, $p_{m}$ is that from a male perspective, $p_{ff}$ is the consanguinity between two random female neighbours, and $p_{mf}$ is that between a random female and a random male neighbour. Given that, for both haploidy and diploidy, there is no sex difference in the consanguinity to self and in the consanguinity to a female patch mate in our model, we may rewrite the above expression as

$(\frac{\partial W_{f}}{\partial x}+\frac{\partial W_{f}}{\partial x^{'}}r)+(\frac{\partial W_{m}}{\partial x}+\frac{\partial W_{m}}{\partial x^{'}}r)$, (S20)

where *r =* $p_{ff}/p_{f}$ *=* $p_{mf}/p_{m}$ is the genetic relatedness between patch mates before dispersal.

Relative fitness:

Under the assumption of vanishingly low variation in traits values in the population (Taylor & Frank, 1996), for both ploidies the fitness of a female (defined by the expected number of offspring she has) can be written as

$w_{f}=(1-ay(1-x^{'}))(\frac{n(1-x)(1-y^{'})}{(1-c)(1-m)n(\bar{x}+(1-\bar{x})\bar{y})(1-a(1-\bar{x})\bar{y})+(1-m)n(1-x^{'})(1-y^{'})(1-a(1-x^{'})y^{'})}+\frac{n(1-c)(x+(1-x)y^{'})}{n(1-m)(1-a(1-\bar{x})\bar{y})(1-c\bar{x}-c(1-\bar{x})\bar{y})})$, (S21)

where *m* is the proportion of offspring that are males. Dividing that by the mean fitness of females (i.e. 1 / (1 – *m*); which is obtained by evaluating eqn S21 at *x = x' = x̄* & *y = y' = ȳ*) gives the relative fitness of a female:

$W_{f}=(1-ay(1-x^{'}))(\frac{(1-x)(1-y^{'})}{(1-c)(\bar{x}+(1-\bar{x})\bar{y})(1-a(1-\bar{x})\bar{y})+(1-x^{'})(1-y^{'})(1-a(1-x^{'})y^{'})}+\frac{(1-c)(x+(1-x)y^{'})}{(1-a(1-\bar{x})\bar{y})(1-c\bar{x}-c(1-\bar{x})\bar{y})})$. (S22)

Then, under the same assumption, for both ploidies the fitness of a male (defined by the expected number of offspring he has) can be written as

$w_{m}=\frac{1-m}{m}(1-a(1-x^{'})y^{'})(\frac{n(1-x^{'})(1-y^{'})}{(1-c)(1-m)n(\bar{x}+(1-\bar{x})\bar{y})(1-a(1-\bar{x})\bar{y})+(1-m)n(1-x^{'})(1-y^{'})(1-a(1-x^{'})y^{'})}+\frac{n(1-c)(x^{'}+(1-x^{'})y^{'})}{n(1-m)(1-a(1-\bar{x})\bar{y})(1-c\bar{x}-c(1-\bar{x})\bar{y})})$. (S23)

Dividing that by the mean fitness of males (i.e. 1 / *m*) gives the relative fitness of a male:

$W_{m}=(1-a(1-x^{'})y^{'})(\frac{(1-x^{'})(1-y^{'})}{(1-c)(\bar{x}+(1-\bar{x})\bar{y})(1-a(1-\bar{x})\bar{y})+(1-x^{'})(1-y^{'})(1-a(1-x^{'})y^{'})}+\frac{(1-c)(x^{'}+(1-x^{'})y^{'})}{(1-a(1-\bar{x})\bar{y})(1-c\bar{x}-c(1-\bar{x})\bar{y})})$, (S24)

which would be identical to that of a female (i.e. $W_{f}$) if all the *x* and *y* of the latter are replaced with *x'* and *y'*. Given that $\partial W_{m}/\partial x$ = 0 and $\partial W_{m}/\partial x^{'}$ = $(\partial W_{f}/\partial x)+(\partial W_{f}/\partial x^{'})$, we can rewrite Expression S20 as

$\frac{\partial W_{f}}{\partial x}(1+r)+\frac{\partial W_{f}}{\partial x^{'}}2r$. (S25)

Relatedness:

Under haploidy, the consanguinity to self ($p_{f}$; Bulmer, 1994) is 1 so that the relatedness of interest equals the consanguinity to a patch mate, which stabilises at

$p_{ff}=s \left( \frac{1}{2}+\frac{1}{2}p_{ff} \right)+\left( 1-s \right)\left( \frac{\left( 1-\bar{x} \right)\left( 1-\bar{y} \right)}{1-c\left( \bar{x}+\left( 1-\bar{x} \right)\bar{y} \right)} \right)^{2}p_{ff}$, (S26)

where *s* is again the probability that two randomly chosen juveniles in a patch have the same mother. That is, if the two females have the same mother, there is a probability of 1/2 that either the maternal or the paternal allele is passed to both. Otherwise, given that the consanguinity between their father and mother is $p_{ff}$, that between the maternal and paternal alleles in the two females is also $p_{ff}$. If they have different mothers, the consanguinity will equal $p_{ff}$ if their mothers are both native and 0 otherwise.

Solving eqn S26 for $p_{ff}$ gives the relatedness of interest under haploidy:

$r=\frac{s}{2(1-(1-s){(\frac{(1-\bar{x})(1-\bar{y})}{1-c(\bar{x}+(1-\bar{x})\bar{y})})}^{2})-s}$. (S27)

Under diploidy, the consanguinity to a patch mate stabilises at

$p_{ff}=s \left( \frac{1}{2} \left( \frac{1}{2}+\frac{1}{2}p_{ff} \right)+\frac{1}{2}p_{ff} \right)+\left( 1-s \right)\left( \frac{\left( 1-\bar{x} \right)\left( 1-\bar{y} \right)}{1-c\left( \bar{x}+\left( 1-\bar{x} \right)\bar{y} \right)} \right)^{2}p_{ff}$. (S28)

That is, if the two females have the same mother, there is a probability of 1/2 that either maternal or paternal alleles are picked twice among them, which then has a probability of 1/2 that they get the same allele of that parent (otherwise, the probability that the two alleles of that parent are identical by descent is $p_{ff}$). If the two alleles picked from the full siblings are not from the same parental origin, then the probability that they are identical by descent is again $p_{ff}$. If the two females have different mothers, the consanguinity between them will be $p_{ff}$ if their mothers are both native and 0 otherwise.

The consanguinity to self for diploidy is

$p_{f}=\frac{1}{2}+\frac{1}{2} p_{ff}$. (S29)

That is, 1 if the same allele from the focal locus is picked twice and $p_{ff}$ (i.e. the consanguinity between her parents) otherwise. Solving eqn S28 for $p_{ff}$ and dividing that by $p_{f}$ gives the relatedness of interest to the evolution of voluntary dispersal, which is identical to the one under haploidy (eqn S27).

Following the same steps above, we find the direction of natural selection acting upon forced dispersal given by

$\frac{\partial W_{f}}{\partial y}(1+r)+\frac{\partial W_{f}}{\partial y^{'}}2r$, (S30)

which depends on the same relatedness and applies to both diploidy and haploidy.

Following the rest of the steps taken in the stability analysis for the clonal model, we recover exact same results (i.e. identical conditions for and stabilised rates under each evolutionary outcome).

Facilitated dispersal

*Fitness function*—Assuming that the mortality cost of modulated dispersal—incurred by the disperser—is lower than that of voluntary dispersal (i.e. dispersing without the modulation), we may change the relative fitness function in the previous section into

$W=\left( 1-\left( 1-x^{'} \right)ay \right)(\left( 1-x \right)\left( 1-y^{'} \right)\frac{1}{\left( 1-\left( 1-x^{'} \right)ay^{'} \right)\left( 1-x^{'} \right)\left( 1-y^{'} \right)+\left( 1-\left( 1-\bar{x} \right)a\bar{y} \right)(\bar{x}\left( 1-c \right)+\left( 1-\bar{x} \right)\bar{y}\left( 1-c\left( 1-f \right) \right))}+(x\left( 1-c \right)+\left( 1-x \right)y^{'}\left( 1-c\left( 1-f \right) \right))\frac{1}{\left( 1-\left( 1-\bar{x} \right)a\bar{y} \right)\left( 1-c\bar{x}-c\left( 1-f \right)\left( 1-\bar{x} \right)\bar{y} \right)})$. (S31)

That is, if the focal individual is not voluntarily dispersing or getting modulated, she will compete with her neighbours that are also not dispersing and with the incoming dispersers that have succeeded either voluntary dispersal (expectedly, proportion 1 – *c* of all voluntary dispersers attempting to enter the focal patch) or modulated dispersal (proportion 1 – *c*(1 – *f*), in which the level of facilitation *f* is the proportion of the mortality cost of dispersal that is alleviated by modulation). If the focal individual is dispersing, then the probability that her dispersal is successful depends on whether she receives modulation.

*Hamilton’s rule & relatedness*—Using the kin-selection methodology of Taylor & Frank (1996), we find that an increase in voluntary dispersal is favoured when

$-\frac{c\left( 1-\bar{y}\left( 1-f \right) \right)}{1-c\left( \bar{x}+\left( 1-f \right)\left( 1-\bar{x} \right)\bar{y} \right)}+\frac{\left( 1-\bar{x} \right)\left( 1-\bar{y} \right)^{2}}{\left( 1-c\left( \bar{x}+\left( 1-f \right)\left( 1-\bar{x} \right)\bar{y} \right) \right)^{2}}r+\frac{a\bar{y}}{1-\left( 1-\bar{x} \right)a\bar{y}}r-\frac{a\bar{y}\left( 1-\bar{x} \right)^{2}\left( 1-\bar{y} \right)^{2}}{\left( 1-\left( 1-\bar{x} \right)a\bar{y} \right)\left( 1-c\left( \bar{x}+\left( 1-f \right)\left( 1-\bar{x} \right)\bar{y} \right) \right)^{2}}r>0$, (S32)

which can then be rearranged into Expression 5 in the main text by multiplying both sides by the reproductive value of an individual before dispersal relative to that of an individual after dispersal (i.e. *v* in the main text).

Using the same approach, we obtain the condition for facilitated dispersal to increase:

$-\frac{a\left( 1-\bar{x} \right)}{1-\left( 1-\bar{x} \right)a\bar{y}}+\frac{a\left( 1-\bar{x} \right)^{3}\left( 1-\bar{y} \right)^{2}}{\left( 1-\left( 1-\bar{x} \right)a\bar{y} \right)\left( 1-c\left( \bar{x}+\left( 1-f \right)\left( 1-\bar{x} \right)\bar{y} \right) \right)^{2}}r-\frac{c\left( 1-f \right)\left( 1-\bar{x} \right)}{1-c\left( \bar{x}+\left( 1-f \right)\left( 1-\bar{x} \right)\bar{y} \right)}r+\frac{\left( 1-\bar{x} \right)^{2}\left( 1-\bar{y} \right)}{\left( 1-c\left( \bar{x}+\left( 1-f \right)\left( 1-\bar{x} \right)\bar{y} \right) \right)^{2}}r>0$, (S33)

which can then be rearranged into Expression 6 in the main text.

In this model, the relatedness between neighbours before dispersal takes place is given by

$r=s+(1-s){(\frac{(1-\bar{x})(1-\bar{y})}{1-c(\bar{x}+\left( 1-f \right)\left( 1-\bar{x} \right)\bar{y})})}^{2}r$. (S34)

Solving that for *r* gives *r = s* / (1 – (1 – *s*) ((1 – *x̄*) (1 – *ȳ*) / (1 – *c*(*x̄* + (1 – *f*)(1 – *x̄*) *ȳ*))) ²), which decreases with a higher level of facilitation. Here, we see that—given that more dispersers are going to successfully reach their destination patch—compared to voluntary dispersal the same level of facilitated dispersal will lead to a lower relatedness. Additionally, we see that—given that, for the same level of dispersal, there are more modulated dispersers reaching the focal patch and thus sharing the reduced competition owing to dispersal—both the second term of Expression S32 and the fourth term of Expression S33 (which reflect the indirect benefit owing to reduced competition among the actor’s neighbours) decrease with a higher level of facilitation. In these two regards, the practice of dispersal facilitation weakens the selection of dispersal.

*Rates & conditions*—Following the steps in the first section of the supplementary material, we obtain the stabilised rate of voluntary dispersal when it is the only form, which is identical to the one yielded in the first model (shown in eqn S9). Substituting this rate of voluntary dispersal and its corresponding stabilised relatedness into Expression S33 and setting *ȳ =* 0, we find the condition that facilitated dispersal is disfavoured to emerge and thus “voluntary dispersal only” is a stable outcome:

$a>\frac{c(2-s(1-f)(1+\sqrt{1+\frac{4c^{2}(1-s)}{s^{2}}}))}{2(1-c)}$. (S35)

This threshold value increases with a lower degree of sibship, a higher cost of dispersal and a higher level of facilitation. If *f =* 0, this threshold value would be identical to the threshold value for forced dispersal to be favoured in the first model (i.e. Expression S10), meaning that the condition for the “voluntary dispersal only” outcome is stricter here.

Different from the one in the forced dispersal model, when *s =* 1 this threshold value here is *cf /* (1 – *c*)—which is always positive. That is, if the cost of modulation times the success rate of voluntary dispersal (i.e. 1 – *c*) is lower than the mortality cost of dispersal that modulation gets to reduce (i.e. *cf*), facilitated dispersal is favoured regardless of local relatedness. Also, here modulated dispersal can be favoured even if *a > c*. In fact, the threshold value would be effectively infinite in the limit of deadly dispersal (i.e. *c* $≐$ 1; but not in the limit of costless modulated dispersal *f* $≐$ 1).

Then, using the same method, we obtain the stabilised rate of facilitated dispersal when it is the only form of dispersal:

$y^{*}=\frac{2s}{2a+(1+a+c(1-f))s+\sqrt{4a^{2}(1+s)+{(1-a+c(1-f))}^{2}s^{2}}}$, (S36)

which increases with a higher degree of sibship, a lower cost of dispersal, a lower cost of modulation and a higher level of facilitation and peaks at the group optimum calculated based on the facilitated cost of dispersal 1 */* (1 + *c*(1 – *f*)) in the limit of costless modulation (*a* $≐$ 0). If *f =* 0, this rate would be identical to the rate of forced dispersal when it is the only form of dispersal (i.e. eqn S11), meaning that this rate—in the same region—is always higher than the forced dispersal rate.

Substituting this rate S36 of facilitated dispersal and its corresponding stabilised relatedness into Expression S32 and setting *x̄ =* 0, we find the condition that voluntary dispersal is disfavoured to emerge and thus “facilitated dispersal only” is a stable outcome. Setting *s =* 1 in this threshold value of the cost of modulation, we obtain a sufficient condition for facilitated dispersal to be the only form of dispersal favoured:

$a<\frac{cf(1-c^{2}(1-f))}{(1-c)(1-c(c(1-f)-2f))}$, (S37)

which increases with a higher cost of dispersal and a higher level of facilitation and is always positive (and, again, becomes effectively infinite in the limit of deadly dispersal). This sufficient condition is a subregion of the parameter space that violates Expression S35.

Then, following the steps in the first section, we obtain the stabilised rate of voluntary dispersal at the intermediate parameter space where both forms of dispersal coexist *x* = F*(*a*, *c*, *s*, *f*), which is a function of all four model parameters and increases with a higher degree of sibship and a lower cost of dispersal. And, the stabilised rate of facilitated dispersal at the region *y* = F*(*a*, *c*, *s*, *f*) is also a function of the four parameters and increases with a lower cost of modulation and a higher level of facilitation. Finally, adding the rate of voluntary dispersal and the rate of facilitated dispersal multiplied by the proportion of individuals that do not voluntarily disperse gives the overall level of dispersal (i.e. *x* +* (1 – *x**) *y**) in the coexistence region.

Given that the rate of facilitated dispersal when it is the only form of dispersal (eqn S36) also increases with a higher degree of sibship and a lower cost of dispersal and that the rate of voluntary dispersal (eqn S9) is always higher than the would-be rate of facilitated dispersal when the former is the only form of dispersal, we know that the adoption of facilitated dispersal will not change the monotonic relationships between the overall rate of dispersal and (1) dispersal cost and (2) sibship. That is, the overall dispersal rate increases with a higher degree of sibship and a lower cost of dispersal. However, different from the model of forced dispersal, here—given that the rate of facilitated dispersal when it is the only form of dispersal may and may not be lower than the would-be rate of voluntary dispersal in that region (see Figure S3)—we know that the overall dispersal rate can increase or decrease with a lower cost of modulation or a higher level of facilitation.

*Inclusive-fitness interests of the recipient*—To reveal the inclusive-fitness interests of the recipient of modulation and thus understand the potential conflicts between the actor and the recipient, we modify the model of facilitated dispersal above. Specifically, we grant the recipients of modulation an option to stay regardless of the modulation at no cost to themselves. This hypothetical control that the recipients have takes values in the interval [0, 1], and by setting that level to zero and investigating when it is favoured to deviate from zero we know the condition that modulation is counter to the inclusive-fitness interests of the recipients.

In this case, the relative fitness function is changed into

$W=(1-(1-x^{'})ay)(\frac{\left( 1-x \right)\left( 1-y^{'}\left( 1-z \right) \right)}{\left( 1-\left( 1-x^{'} \right)ay^{'} \right)\left( 1-x^{'} \right)\left( 1-y^{'}\left( 1-z^{'} \right) \right)+\left( 1-\left( 1-\bar{x} \right)a\bar{y} \right)\left( \bar{x}\left( 1-c \right)+\left( 1-\bar{x} \right)\bar{y}\left( 1-\bar{z} \right)\left( 1-c(1-f \right)) \right)}+\frac{x(1-c)+(1-x)y^{'}(1-z)(1-c(1-f))}{(1-(1-\bar{x})a\bar{y})(1-c(\bar{x}+(1-f)(1-\bar{x})\bar{y}(1-\bar{z})))})$, (S38)

in which *z* is the probability that the focal individual chooses to stay when she receives modulation, *z'* is the average probability that a neighbour of her chooses to stay when the neighbour receives modulation, and *z̄* is the average probability that a recipient chooses to stay among the whole population. Setting all these probabilities to zero (*z =* 0; *z' =* 0; *z̄ =* 0) recovers the relative fitness function in the facilitated dispersal model (eqn S31).

Using the kin selection methodology of Taylor & Frank (1996), we obtain the marginal fitness of the staying probability:

$\frac{c(1-f)(1-\bar{x})\bar{y}}{1-c(\bar{x}+\left( 1-f \right)\left( 1-\bar{x} \right)\bar{y}\left( 1-\bar{z} \right))}-\frac{\left( 1-\bar{x} \right)^{2}\bar{y}\left( 1-\bar{y}\left( 1-\bar{z} \right) \right)}{\left( 1-c(\bar{x}+\left( 1-f \right)\left( 1-\bar{x} \right)\bar{y}\left( 1-\bar{z} \right)) \right)^{2}}r$. (S39)

Specifying the relationship between relatedness and the two forms of dispersal (see eqn S34) and setting *z̄ =* 0, we find the condition that the probability is favoured to deviate from zero, which implies that the recipient’s inclusive-fitness interests are violated:

$c(1-f)-\frac{(1-\bar{x})(1-\bar{y})}{1-c(\bar{x}+\left( 1-f \right)\left( 1-\bar{x} \right)\bar{y})}\frac{s}{1-(1-s){(\frac{\left( 1-\bar{x} \right)\left( 1-\bar{y} \right)}{1-c(\bar{x}+\left( 1-f \right)\left( 1-\bar{x} \right)\bar{y})})}^{2}}>0$. (S40)

Setting *f* = 0 & *x̄ =* 0, we find that, when modulated dispersal incurs the same dispersal cost and is the only form of dispersal, the substituted Expression S40 is fulfilled if its rate exceeds the would-be voluntary dispersal rate (eqn S9). Given that, in the parameter space where non-facilitating modulated dispersal is the only form of dispersal favoured, its stabilised rate (eqn S11) is always higher than that, modulation must be against the recipient’s inclusive-fitness interests (i.e. forced dispersal) in the region. Setting *f* = 0 and substituting the stabilised rates of modulated dispersal and voluntary dispersal in the coexistence region of both forms of dispersal into Expression S40, we find that the substituted Expression S40 is always fulfilled, meaning that in the coexistence region modulated dispersal which does not improve dispersal success is, again, against the recipient’s inclusive-fitness interests (i.e. forced dispersal).

Then, leaving *f* unspecified and setting *x̄ =* 0, we find that, when modulated dispersal incurs a reduced dispersal cost and is the only form of dispersal, Expression S40 is fulfilled when its rate exceeds the would-be rate of voluntary dispersal calculated based on the facilitated dispersal cost (i.e. eqn S9 with *c* replaced by *c* (1 – *f*)). This threshold value of the rate of modulated dispersal is always higher than that when modulated dispersal does not improve dispersal success and increases with a higher level of facilitation, a higher degree of sibship and a lower cost of dispersal. Given that, when facilitated dispersal is the only form of dispersal favoured, its stabilised rate (eqn S36) can be lower than eqn S9, we know that facilitated dispersal need not be unwelcome. Specifically, comparing eqn S36 to the threshold value of the rate of modulated dispersal, we find that the cost of modulation must be lower than a point, which is a function of the cost of dispersal, the level of facilitation and the degree of sibship (and equals 0 when *s =* 1), so that its favoured rate can exceed that threshold value and thus cause modulation to be unwelcome. Similarly, substituting the stabilised rates of voluntary dispersal and facilitated dispersal in the coexistence region into Expression S40, we find that the condition may and may not be fulfilled in the region. In general, when *a > c* (1 – *f*), the favoured rate of modulated dispersal will not fulfil Expression S40 and thus modulated dispersal will not be unwelcome.

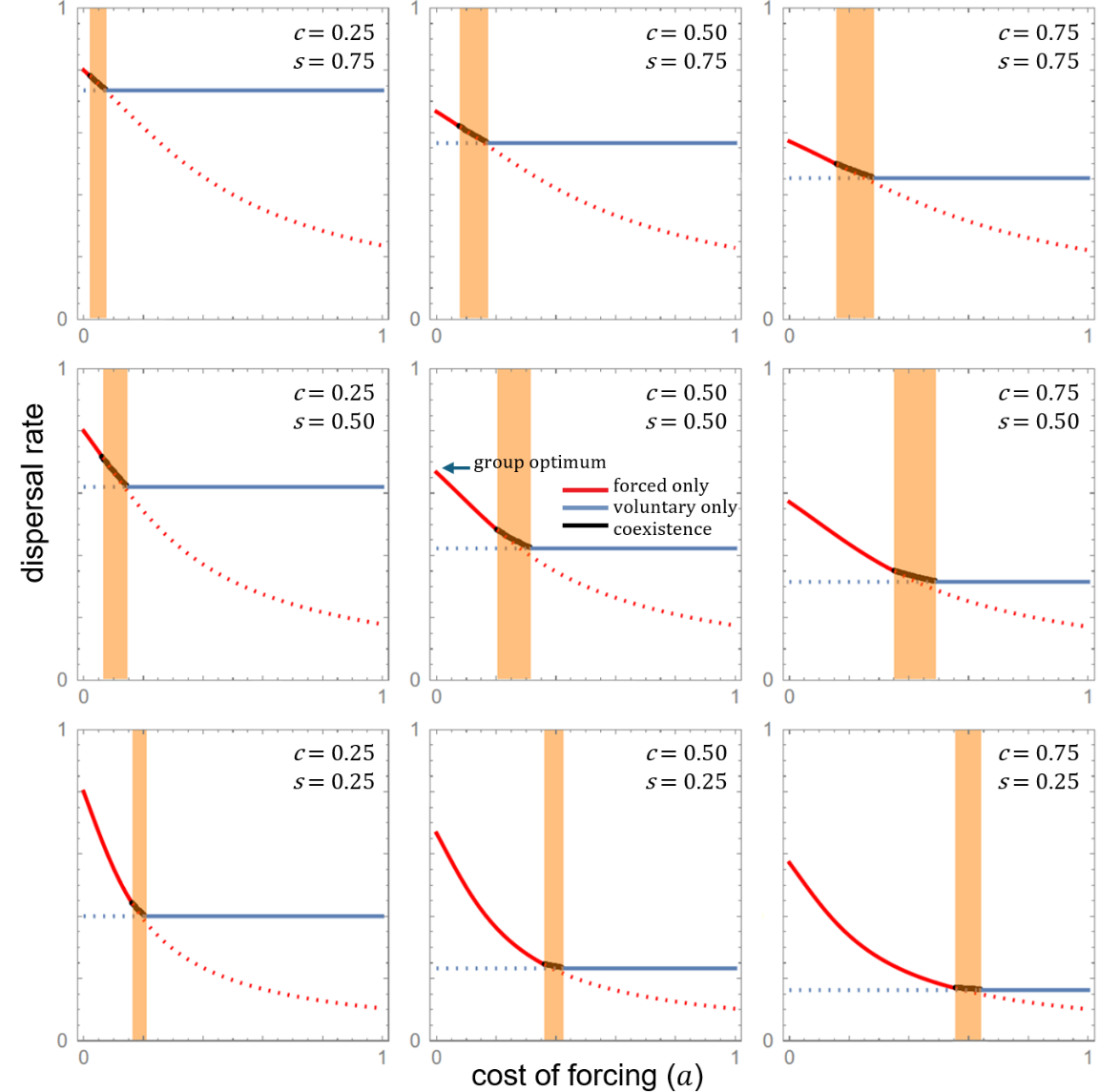

**Figure S1 | Three stable outcomes and their rates, when social modulation does not improve dispersal success.** Both the modes and the overall level of dispersal depend on an interplay of the cost of dispersal (*c*), the degree of sibship (*s*) and the cost of forcing (*a*). The red solid line is the stabilised dispersal rate when only forced dispersal is favoured and the blue solid line is that when only voluntary dispersal is favoured. The black line covers the region where both forms are favoured at an intermediate rate. For comparison, the blue dotted line is the would-be voluntary dispersal rate if forcing is nonexistent while the red dotted line is the would-be forced dispersal rate if no individual voluntarily disperses. The point where the red line touches the vertical axis corresponds to the group optimum of dispersal.

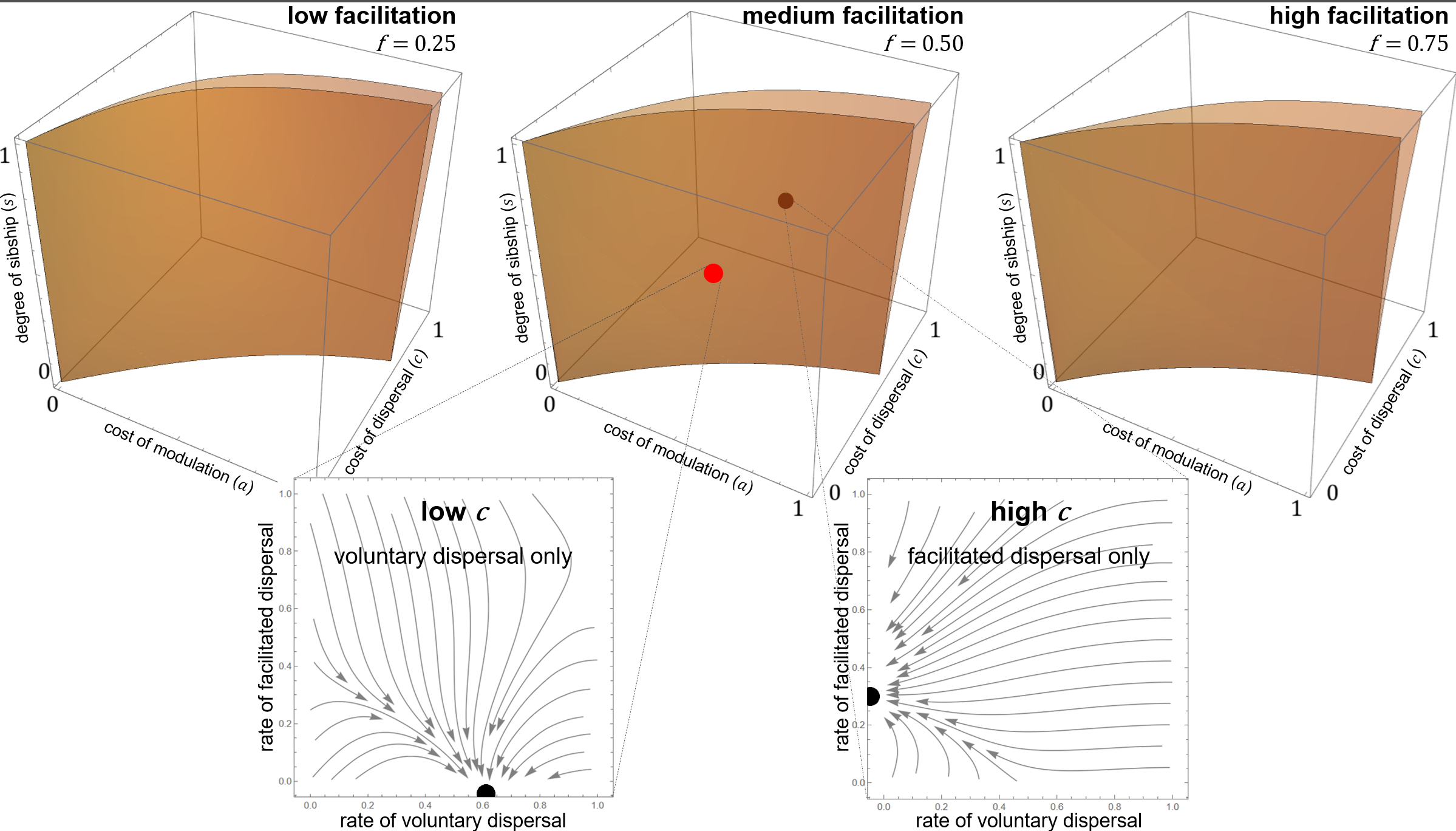
 **Figure S2 | Three stable outcomes, when social modulation improves dispersal success.** For high level of facilitation (high *f*), low cost of modulation (low *a*), high cost of dispersal (high *c*) and low degree of sibship (low *s*), only facilitated dispersal obtains (i.e. portion of parameter space lying behind the orange volume). For low level of facilitation (low *f*), high cost of modulation (high *a*), low cost of dispersal (low *c*) and high degree of sibship (high *s*), only voluntary dispersal obtains (i.e. portion of parameter space lying in front of the orange volume). For intermediate level of facilitation (medium *f*), intermediate cost of modulation (medium *a*), intermediate cost of dispersal (medium *c*) and intermediate degree of sibship (medium *s*), both voluntary and facilitated dispersal coexist (i.e. portion of parameter space lying within the orange volume).

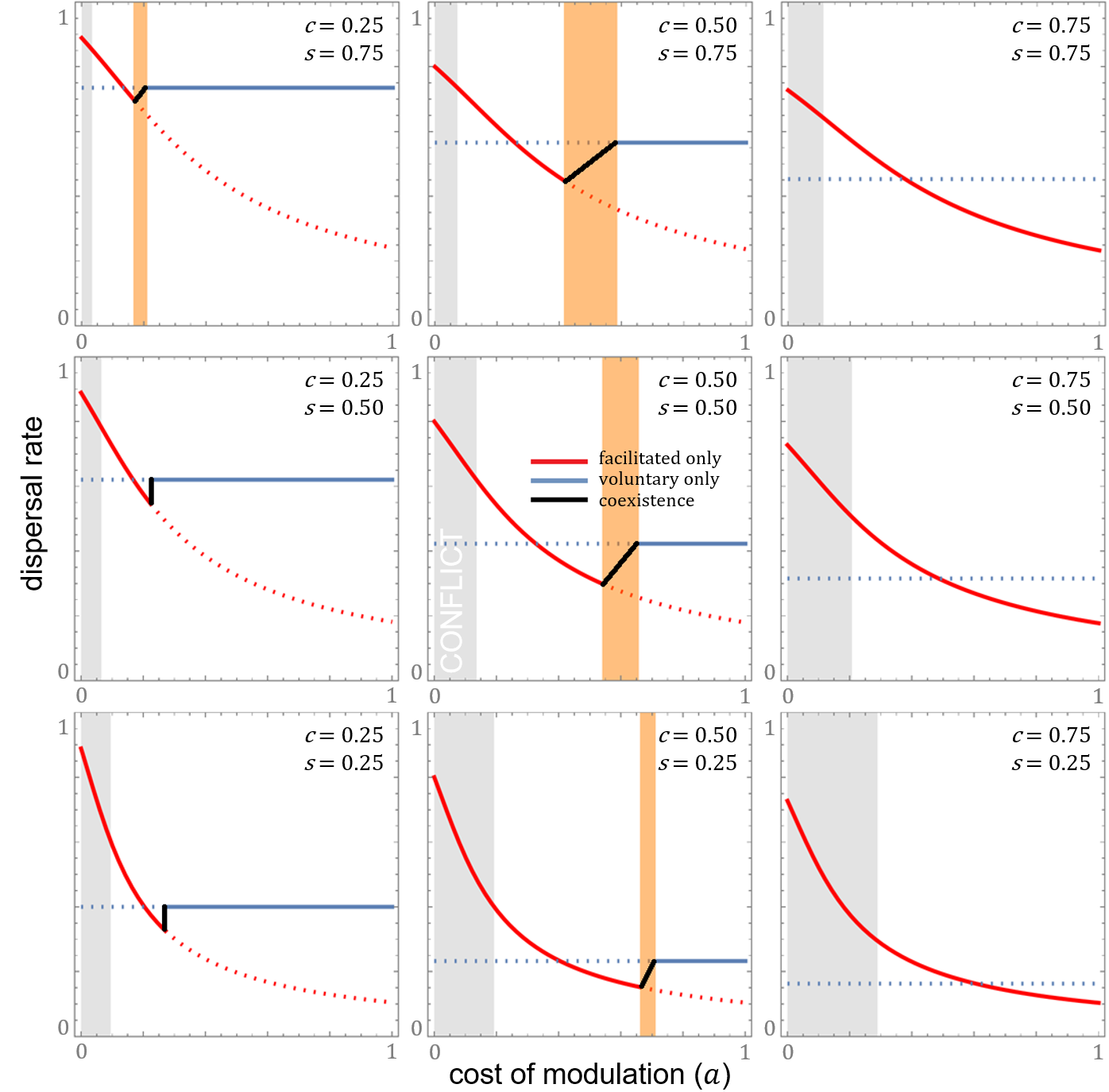

**Figure S3 | Rates of dispersal, when social modulation improves dispersal success.** Although facilitated dispersal always leads to increased willingness to disperse and thus a potential for more dispersal as the cost of modulation reduces, it could reduce the overall dispersal rate favoured in a population. The red solid line is the stabilised dispersal rate when only facilitated dispersal is favoured and the blue solid line is that when only voluntary dispersal is favoured. The black line covers the region that both forms are favoured. For comparison, the blue dotted line is the would-be voluntary dispersal rate if modulation is nonexistent while the red dotted line is the would-be facilitated dispersal rate if no one disperses by themselves. The grey region is where modulation acts against the recipients’ inclusive-fitness interests. Here, facilitated dispersal is assumed to reduce the cost of dispersal by half (i.e. *f =* 0.5).
